# Dendrimer Delivered shRNA Targeting the CCL20–CCR6 Axis Suppresses Complement-Mediated Microglial Synaptic Pruning and Ameliorates Chronic Neuroinflammation After Repetitive Traumatic Brain Injury

**DOI:** 10.64898/2026.09.04.748639

**Authors:** Karthick Mayilsamy, Parthvi Bharatkumar Patel, Ryan Green, Sashank Bikkasani, Kristina Tosi, Shreyasi Majumdar, Santosh Kumar Prajapati, Eleni Markoutsa, Tiara Wolf, Jennifer Guergues, Stanley M. Stevens, Alison Willing, Shyam Mohapatra, Subhra Mohapatra

## Abstract

Repetitive traumatic brain injury (rTBI) induces persistent microglial activation and chronic neuroinflammation, yet the upstream signals driving long⍰term synaptic injury remain unclear. In this study, we identify the CCL20–CCR6 chemokine axis as a critical regulator of sustained microglial activation and complement⍰dependent synaptic loss after rTBI. Proteomic profiling at 30 days post⍰injury (dpi) showed broad normalization of complement⍰linked inflammatory and synaptic pathways in the cortex and hippocampus, underscoring a mechanistic link between chemokine signaling, microglial activation, and synaptic vulnerability. To therapeutically target this axis, we developed a dendrimer⍰based shRNA platform (shCombo⍰DPX) that simultaneously silences CCL20 and CCR6. Intranasal and intravenous delivery in rTBI mice effectively reduced CCL20–CCR6 expression, attenuated chronic microgliosis and astrogliosis, and suppressed complement activation. Treatment limited microglial synaptic engulfment, preserved synaptic proteins, restored BDNF levels, and improved motor, anxiety⍰related, and cognitive outcomes. In microglia–neuron coculture systems, CCL20 silencing reduced LPS⍰induced complement signaling and prevented synaptic loss, neuronal apoptosis, and BDNF depletion. Conversely, exposure to recombinant CCL20 induced dendritic degeneration, caspase⍰3 activation, microglial reactivity, complement dysregulation, and synaptic injury both in vitro and in vivo. Collectively, these findings establish CCL20–CCR6 as a key upstream driver of chronic complement⍰mediated synaptic degeneration after rTBI and support dendrimer⍰delivered shRNA therapy as a targeted strategy to mitigate long⍰term neurodegeneration.

## INTRODUCTION

Repeated traumatic brain injury (rTBI) has become a significant neurological health concern, particularly among athletes, military personnel, and individuals exposed to recurrent concussive events. Unlike a single mild TBI, which often resolves clinically within days to weeks, repeated injuries occurring within the vulnerable recovery window can exacerbate neuropathology and result in cumulative, long-lasting neurodegenerative consequences [1, 2]. Both clinical and experimental studies demonstrate that rTBI increases the risk of persistent mood disturbances, cognitive impairment, and neuropsychiatric symptoms, while elevating susceptibility to chronic neurodegenerative diseases, including chronic traumatic encephalopathy (CTE) and Alzheimer’s disease (AD) [3, 4]. Recurrent injuries can initiate a chronic, evolving pathophysiological state driven by sustained neuroinflammation and progressive neurodegeneration. As the secondary injury cascade unfolds, the brain transitions into a prolonged inflammatory environment that promotes a self-perpetuating cycle of inflammation and neurodegeneration underlying the long-term burden of rTBI [5, 6].

Microglial reactivity is a hallmark of TBI, and microgliosis can persist for years following the initial insult [7–9]. Although acute microglial activation can support tissue repair and restore homeostasis, chronically reactive microglia produce proinflammatory and cytotoxic mediators that exacerbate neuronal damage [10–12]. In the context of rTBI, each impact may trigger another wave of microglial activation, creating a chronically primed state. This persistent reactivity drives excessive synaptic loss, disrupts neuronal plasticity, and destabilizes network connectivity, supporting the concept that microglial activation is a central driver of cognitive decline that follows rTBI [13, 14].

Traumatic and inflammatory insults strongly activate the complement system through reactive glial cells, driving sustained neuroinflammation and chronic pathology[15–17]. While complement activity aids debris clearance and early repair, prolonged activation becomes maladaptive, amplifying inflammation and promoting neurodegeneration. C1q deposition on stressed synapses initiates the classical pathway and downstream signaling [18], producing opsonins (C3b/iC3b) and anaphylatoxins (C3a, C5a) that heighten glial reactivity, recruit peripheral immune cells, and enhance phagocytosis. Accumulation of C1q and C3 at vulnerable synapses marks them for microglial engulfment, leading to long-term synaptic loss [19, 20]. Effectors such as C5a further sustain inflammation through chemotaxis and cytokine release[21]. Chronic complement activation is associated with progressive synapse elimination, dendritic spine loss, and memory deficits. Consequently, targeting complement components has emerged as a therapeutic strategy to limit chronic neuroinflammation and preserve synaptic integrity after rTBI [22, 23].

C⍰C motif chemokine ligand 20 (CCL20), which signals exclusively through its lone receptor CCR6, plays a sustained and amplifying role in chronic neuroinflammation after TBI. CCL20 functions both as a chemoattractant and as a potent activator of glial cells [24, 25]. Across multiple CNS injury models, CCL20 is upregulated in neurons, astrocytes, and microglia under inflammatory stress, where it recruits Th17 cells and other leukocytes and helps maintain a prolonged cytokine⍰rich environment [24, 26, 27]. Our previous studies demonstrated that CCL20 contributes to TBI⍰related pathology, including neurodegeneration, microgliosis, astrogliosis, and retinal injury [28, 29]. However, whether CCL20 drives chronic microglial activation in a manner that increases neuronal synaptic vulnerability is poorly understood. These findings highlight an important role for CCL20–CCR6 signaling in TBI⍰associated neuroinflammation, although its contribution to synaptic regulation following rTBI remains unclear.

Building on our previous work, we employed an rTBI model to investigate the role of the CCL20–CCR6 axis in chronic synaptic injury and evaluated a dendrimer-based therapeutic strategy to attenuate this signaling pathway. Dendriplex (DPX) therapy carrying plasmids encoding shRNAs targeting both CCL20 and CCR6 produced the most pronounced suppression of neuroinflammation compared with treatment using either individual shRNA alone [29]. Notably, Poly(amidoamine) (PAMAM) dendrimers selectively accumulate in activated microglia and injured neurons, enabling pathology-driven targeting without the need for additional targeting ligands [30, 31]. This intrinsic tropism, together with the proton-sponge effect, facilitates endosomal escape and enhances intracellular gene-silencing efficiency [32]. Their nanoscale size and multivalent surface characteristics further support blood–brain barrier (BBB) penetration through adsorptive-mediated transcytosis and inflammation-enhanced transport mechanisms [33, 34].

Herein, we demonstrate that in a mouse rTBI model, sustained increase in CCL20 promotes prolonged microglial activation and enhanced complement engagement, creating conditions that accelerate synaptic loss. Treatment with shCCL20–CCR6 dendriplex (DPX) reduced CCL20 and CCR6 expression, dampened neuroinflammatory signaling, and limited complement⍰mediated synaptic pruning. These findings identify the CCL20–CCR6 pathway as a key regulator of the inflammatory–complement cascade and suggest that its inhibition may represent a promising approach to mitigate synaptic damage after rTBI.

## RESULTS

### Proteomic changes in the cortex and hippocampus of rTBI mice at 30 dpi

Our previous studies identified CCL20 as a critical neuroinflammatory mediator that contributes to the pathological and behavioral deficits associated with TBI [29, 35]. Building on these findings, the present study investigates the mechanisms by which CCL20 amplifies neuroinflammatory sequelae through its interactions with downstream signaling partners and associated proteins, thereby promoting chronic inflammatory responses following rTBI. To address this, Cortical and Hippocampal tissues were dissected from sham and rTBI mice, and total protein expression was analyzed using an untargeted bottom⍰up mass spectrometry approach. Comparative analysis identified 738 differentially expressed proteins (DEPs) in the cortex and 258 DEPs in the hippocampus between rTBI and sham groups. To visualize global proteomic differences between groups, volcano plots were generated using fold-change and statistical significance analyses (Fig. 1A, B). Proteins were considered significantly differentially expressed using a threshold of log₂ (fold change) ≥ 1 and an adjusted p⍰value ≤ 0.05. Significantly upregulated proteins are shown on the right side of the volcano plots, whereas significantly downregulated proteins are shown on the left (Fig. 1A, B). Corresponding heat maps illustrating the expression patterns of differentially expressed proteins are presented in Fig. S1 A (Cortex) and S1 B (Hippocampus). Pathway analysis was performed separately for each region using Ingenuity Pathway Analysis (IPA, Qiagen), and Gene Ontology functional enrichment and interactome analyses were used to compare protein expression signatures between the two regions (Metascape). A circos plot was used to visualize the similarity in protein expression between cortex and hippocampus (Fig 1C). In the cortex, pathway analysis revealed activation of mitochondrial dysfunction, CLEAR (Coordinated Lysosomal Expression and Regulation) signaling, and granzyme A signaling, whereas vesicle transport, neurotransmitter release, and synaptogenesis were predicted to be suppressed (Fig. 1D, S2 A). In the hippocampus, the top activated pathways included microautophagy, the RHO GTPase cycle, CLEAR signaling, and KEAP1–NRF2 signaling, while RHOGDI signaling and pyrimidine salvage were predicted to be downregulated. IPA was also used to assess predicted disease and functional outcomes associated with the DEP profiles. In the hippocampus, cell migration and immune responses were predicted to be activated, while ubiquitination and hypoplasia were predicted to be inhibited (Fig. 1D, S 2A). Further in the cortex, necrosis, seizures, and movement⍰related disorders were predicted to be activated, whereas neuritogenesis, synaptic processes, and cognition were predicted to be inhibited. To examine CCL20-related chemokine mechanisms, DEP datasets were queried for proteins associated with CCL20 signaling and complement activation using IPA (Fig. 1E, F). Based on the observed protein expression patterns, CCL20 signaling was predicted to be activated in both the hippocampus and cortex. Complement components C3 and C5 were also predicted to be activated in both regions, while C1QC and C4a/C4b showed increased abundance specifically in the hippocampus (1.47⍰fold and 2.24⍰fold, respectively). DEPs known to interact with CCL20 signaling, as well as those linking CCL20 to complement proteins, are shown for both regions (Fig. 1E, F, Table S1, S2). Gene ontology functional enrichment analysis was performed on both datasets, revealing acute inflammatory response, post-translational protein phosphorylation, and neutrophil degranulation among the top-enriched GO terms shared between the brain regions. (Fig 1G, S 2B, S 2C). Overall, the proteomic analysis reveals widespread molecular alterations in the cortex and hippocampus 30 days after rTBI, including significant up⍰ and down⍰regulation of proteins involved in inflammation, synaptic function, and cellular stress responses.

**Figure 1.**
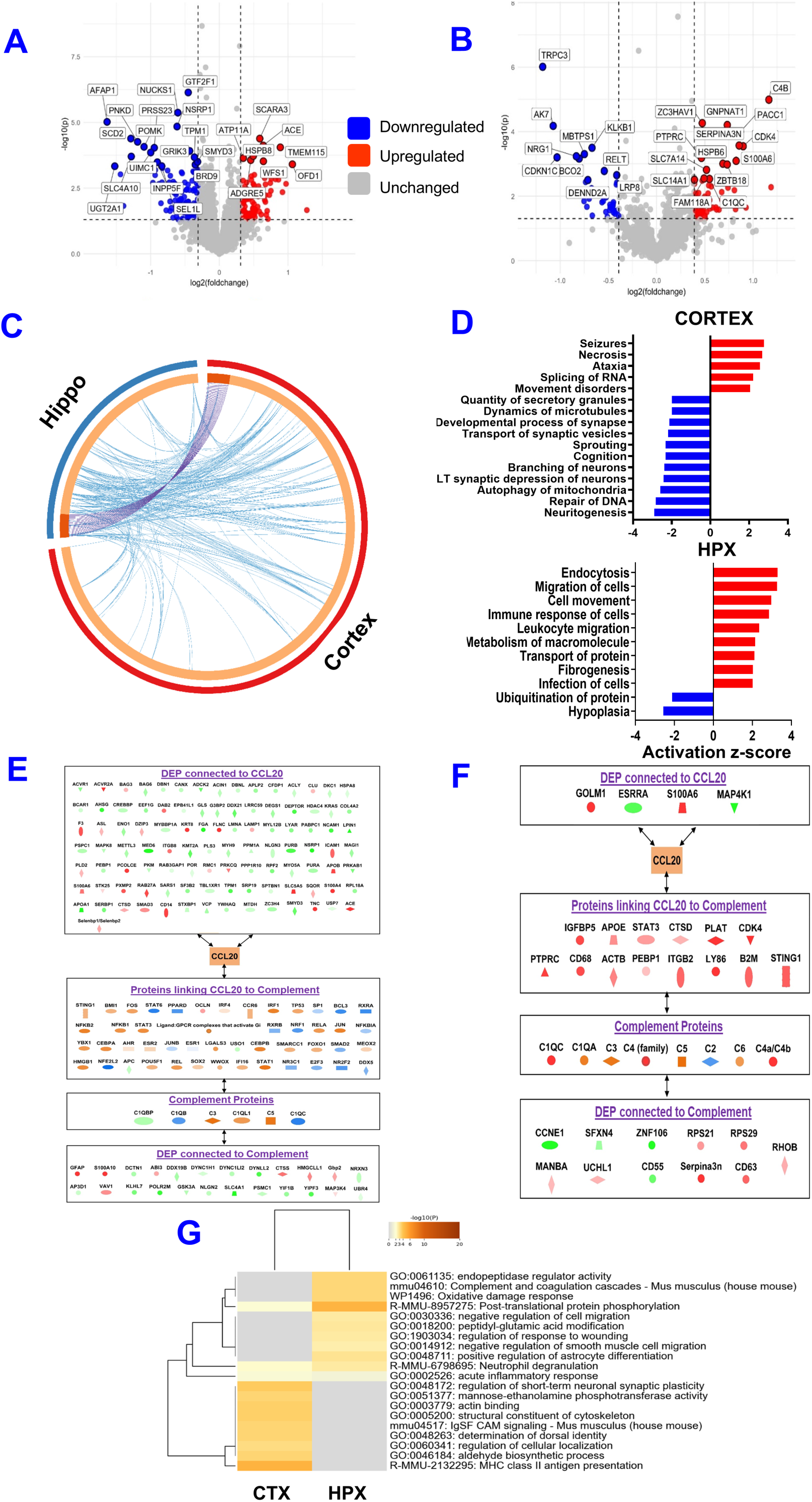
Proteomic alterations in the cortex and hippocampus 30 days after rTBI. Differentially abundant proteins were determined by MSqRob p-value < 0.05, log2FC > |bootstrapping-determined value|, n=3 per group. Volcano plots showing differentially expressed proteins in the cortex **(A)** and hippocampus **(B)** at 30 dpi, highlighting proteins significantly upregulated or downregulated relative to sham controls. Corresponding circos plot of differentially expressed proteins in the cortex (red segment) and hippocampus (blue segment). Proteins that are members of both datasets are linked with purple lines. Proteins that are different in each dataset but fall under the same ontology term are linked with blue lines. **(C)** Histograms summarizing disease⍰ and function⍰associated categories enriched among differentially expressed proteins in the cortex and hippocampus **(D)**, reflecting early molecular signatures of neuroinflammation, synaptic dysfunction, and neuronal stress responses following repeated injury. Ingenuity Pathway Analysis (IPA) illustrating observed and predicted CCL20⍰centered interaction networks in the cortex **(E)** and hippocampus **(F).** Heatmap depicting hierarchical clustering of the top 20 significantly enriched GO terms as ranked by p-value **(G).**

### Microglia as central mediators of CCL20 signaling, complement activation, and synaptic engulfment in rTBI

Dual⍰label immunofluorescence revealed that CCL20 expression colocalizes predominantly with IBA1⁺ microglia, identifying activated microglia as the primary CCL20⍰producing cell population following rTBI. Minimal CCL20 colocalization was observed with GFAP⁺ astrocytes or NeuN⁺ neurons in the cortical injury site, indicating limited chemokine production by these cell types. Although astrocytes exhibited robust GFAP reactivity following rTBI, CCL20 staining remained sparse. Neurons similarly showed minimal CCL20 labeling (Fig. 2A, S3 A-C). Quantitative analyses demonstrated a clear rightward shift in fluorescence intensity distributions in rTBI tissue, reflecting robust microglial upregulation of CCL20 (Fig. 2B). To further validate the cellular source of CCL20 within the chronic neuroinflammatory milieu induced by rTBI, we analyzed *Ccl20* gene expression in isolated microglia by qPCR. The results demonstrated a significant upregulation of *Ccl20* expression in microglia from rTBI brains compared with sham controls. These findings confirm that microglia are a major source of CCL20 in the injured cortex and support their role in sustaining chronic neuroinflammation (Fig. S4 A, B).

**Figure 2.**
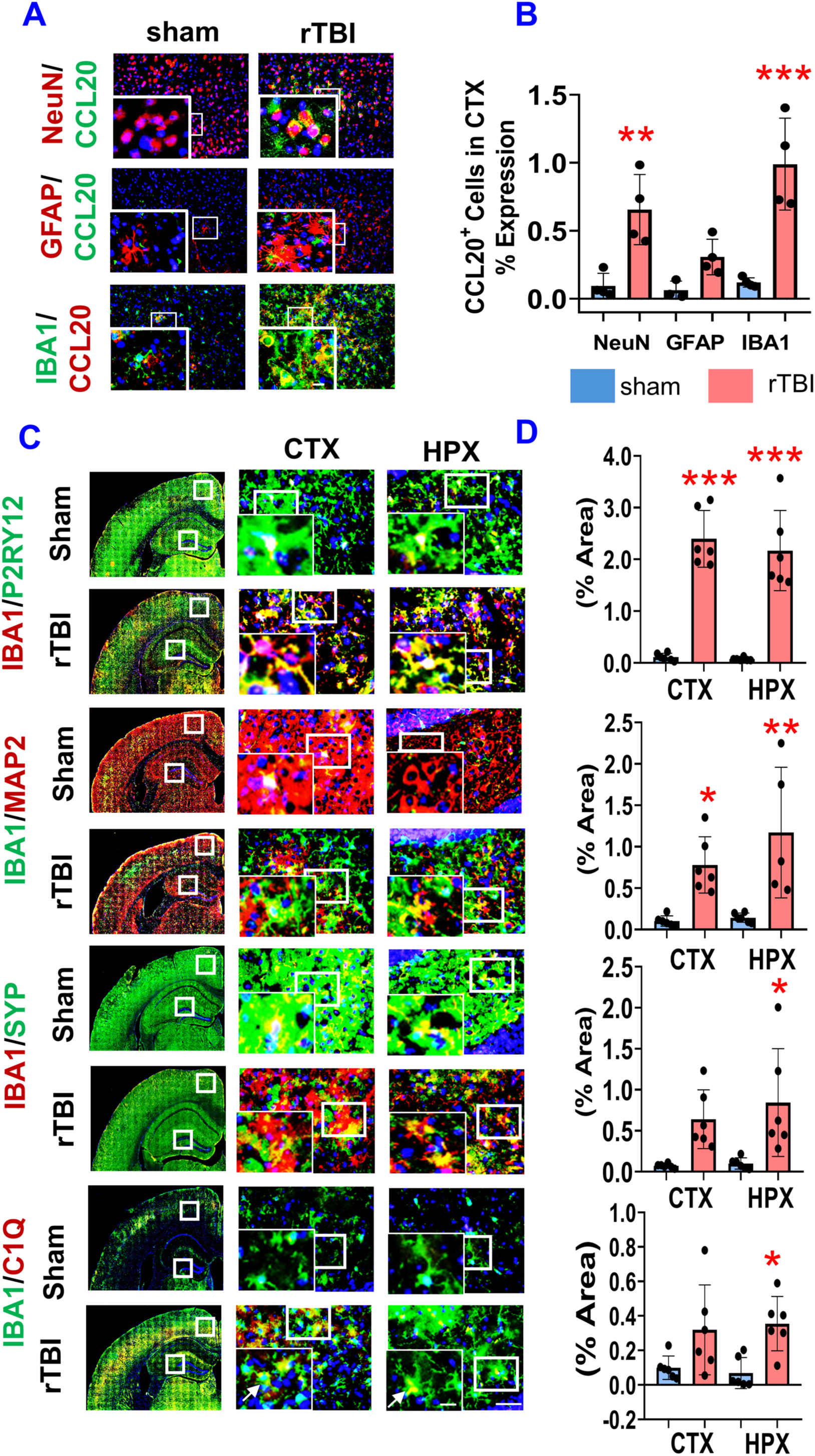
Microglia as central mediators of CCL20 signaling, complement activation, and synaptic engulfment in rTBI. **(A)** Immunostaining of CCL20 expression in neurons (NeuN⁺), astrocytes (GFAP⁺), and microglia (IBA1⁺) within the cortical injury site at 30□ dpi. **(B)** Quantification of CCL20⁺ cells expressed as the percentage of each brain cell type in the cortex (n □=□ 4). **(C)** Representative immunofluorescence images showing double immunostaining of P2RY12 (Green) / IBA1 (Red); IBA1 (Green) / MAP2 (Red); SYP (Green) / IBA1 (Red); IBA1 (Green) and C1Q (Red), counterstained with DAPI in the cortex (CTX) and hippocampus (HPX) at 30 DPI **(D)** Quantification of mean fluorescence intensity ratios (IBA1/P2RY12, IBA1/MAP2, IBA1/SYP) and IBA1⁺/C1q⁺ cells (%Area); n □=□ 6/group. Scale bar: 50 □µm (inset: 10□µm). Data are presented as mean □±□ SD. Statistical analysis: one⍰way ANOVA with Holm–Sidak test; *p□< □0.05, **p □<□ 0.01, ***p □<□ 0.001 vs. sham; #p□ <□ 0.05, ##p □< □0.01 vs. treated groups.

We next examined whether microglia undergo a transition from a homeostatic to a chronically activated state following rTBI by assessing the expression of the homeostatic microglial marker P2RY12 and activation marker IBA1 at 30 dpi (Fig. 2 C, D, S5). Although these markers were selected to distinguish resting from phagocytic microglia, this separation was not feasible because most P2RY12⍰positive cells in sham tissue also expressed IBA1. However, higher⍰magnification imaging showed a clear increase in both the number and intensity of IBA1⍰positive microglia in rTBI mice compared with sham controls. P2RY12 remained detectable but showed reduced signal within IBA1⍰positive cells in rTBI tissue, consistent with a transition toward a reactive phenotype. This activated microglial state persisted up to 120 dpi (Fig. S6 A, B, S7). To visualize microglia–neuron interactions, particularly microglial engagement with dendritic structures, IBA1/MAP2 immunostaining was performed. MAP2 immunoreactivity was markedly reduced in the cortex and hippocampus of rTBI mice at 30 dpi, coinciding with increased IBA1 activation (Fig. 2C, D, S5). By 120 dpi, MAP2 loss and microglial activation remained evident, though microglial reactivity was lower than at 30 dpi (Fig. S6 A, B, S7). To evaluate microglia-mediated synaptic engulfment, synaptophysin (SYP) and IBA1 co⍰expression were assessed. Sham mice showed uniform synaptophysin puncta in both the cortex and hippocampal CA1 region. In contrast, rTBI mice exhibited marked synaptophysin loss at 30 dpi, particularly in regions with high IBA1 density, with SYP proteins frequently clustered within microglial accumulations (Fig. 2 C, D, S5). To examine complement involvement in synaptic damage, IBA1/C1q co⍰staining was performed. Activated microglia showed increased co⍰expression with C1q in both the cortex and hippocampus at 30 dpi compared with sham mice (Fig. 2 C, D, S5). Elevated IBA1–C1q overlap persisted through 120 dpi, albeit at lower levels than at 30 dpi (Fig. S6 A, B, S7).

**Figure 3.**
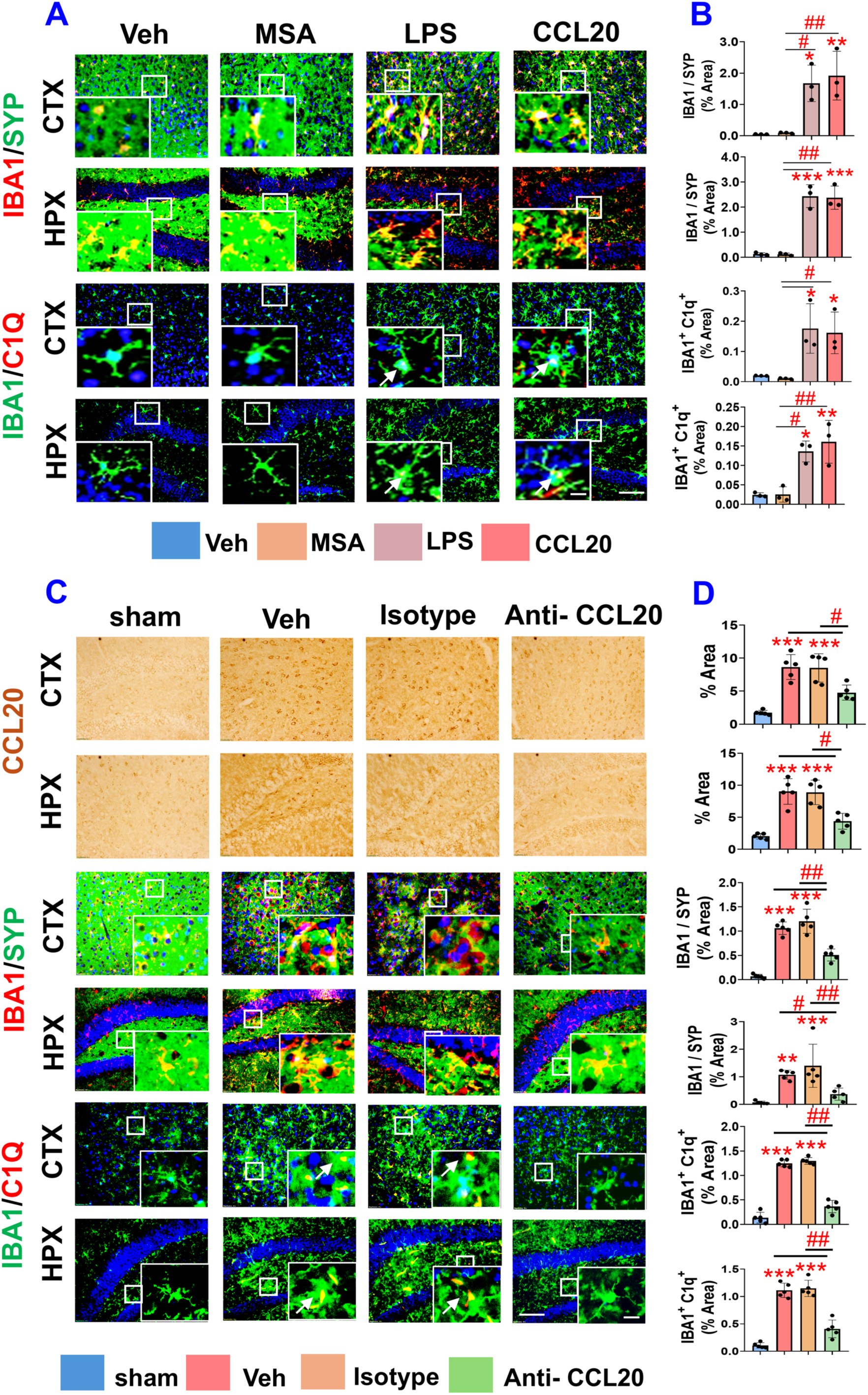
CCL20 signaling drives microglial activation and serves as a therapeutic target for complement⍰mediated synaptic damage after rTBI. (**A–B**) Microglial activation and synaptic loss following administration of recombinant CCL20. (**A**) Representative immunofluorescence images showing double labeling of SYP (Green) / IBA1 (Red); IBA1 (Green) / C1Q (Red); counterstained with DAPI in cortex (CTX) and hippocampus (HPX) at 14 □days post⍰injection with vehicle, mouse serum albumin (MSA), lipopolysaccharide (LPS), or recombinant CCL20. (**B**) Quantification of mean fluorescence intensity ratios (IBA1/SYP) and IBA1⁺/C1q⁺ cells (%Area); n □=□ 3/group. (C-D) Administration of a CCL20⍰neutralizing antibody attenuates synaptic damage in rTBI mice. (C) Representative brightfield images depicting CCL20 expression and immunofluorescence images showing SYP (Green) / IBA1 (Red); IBA1 (Green) / C1Q (Red); counterstained with DAPI in CTX and HPX at 30□ dpi following treatment with vehicle, Isotype Ig, or CCL20⍰neutralizing antibody. (D) Quantification of CCL20 expression (%Area), mean fluorescence intensity ratio (IBA1/SYP) and IBA1⁺/C1q⁺ cells (% area); n□ =□ 5/group. Scale bar: 50 □µm (inset: 10 □µm). Data are presented as mean□ ±□ SD. Statistical analysis: one⍰way ANOVA with Holm–Sidak test; *p□ <□ 0.05, **p □<□ 0.01, ***p□ <□ 0.001 vs. sham; #p□ <□ 0.05, ##p □<□ 0.01 vs. treated groups.

**Figure 4.**
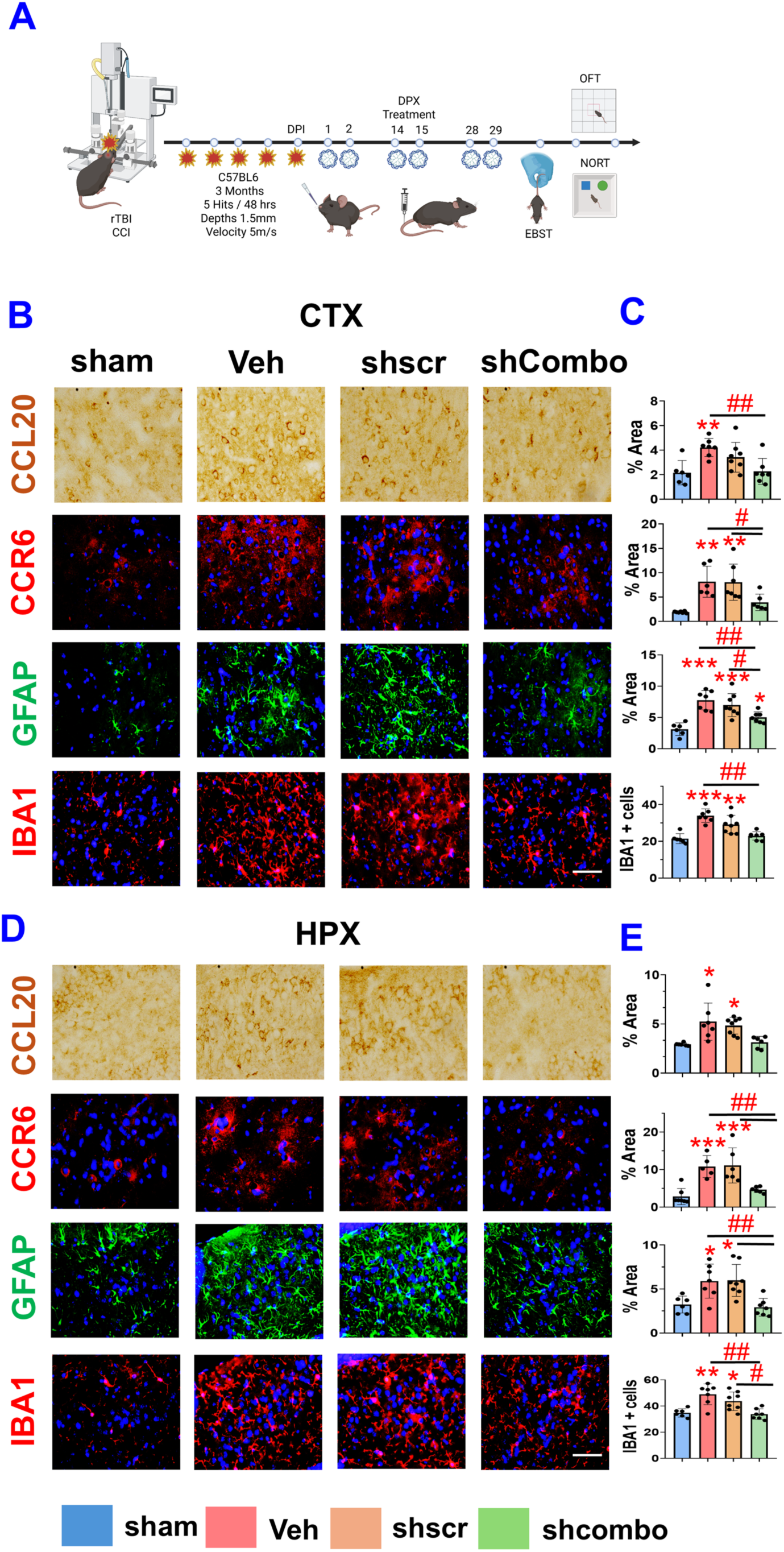
Treatment with shCombo⍰DPX mitigates glial activation and attenuates synaptic damage in rTBI mice. (A–E) Treatment with shCombo⍰DPX improves rTBI pathology. **(A)** Experimental design for shCombo⍰DPX administration in rTBI mice. **(B–E)** Representative brightfield and fluorescent images of cortex (CTX, B) and hippocampus (HPX, D) sections stained for CCL20, CCR6, GFAP, and IBA1 at 30 □dpi, with corresponding quantification for cortex **(C)** and hippocampus **(E)**. Scale bar: 50 □µm (inset: 10 □µm). n□ =□ 6/group; Data are presented as mean□ ±□ SD. Statistical analysis: one⍰way ANOVA with Holm–Sidak test; *p □<□ 0.05, **p □<□ 0.01, ***p□ <□ 0.001 vs. sham; #p □< □0.05, ##p □<□ 0.01 vs. treated groups.

**Figure 5.**
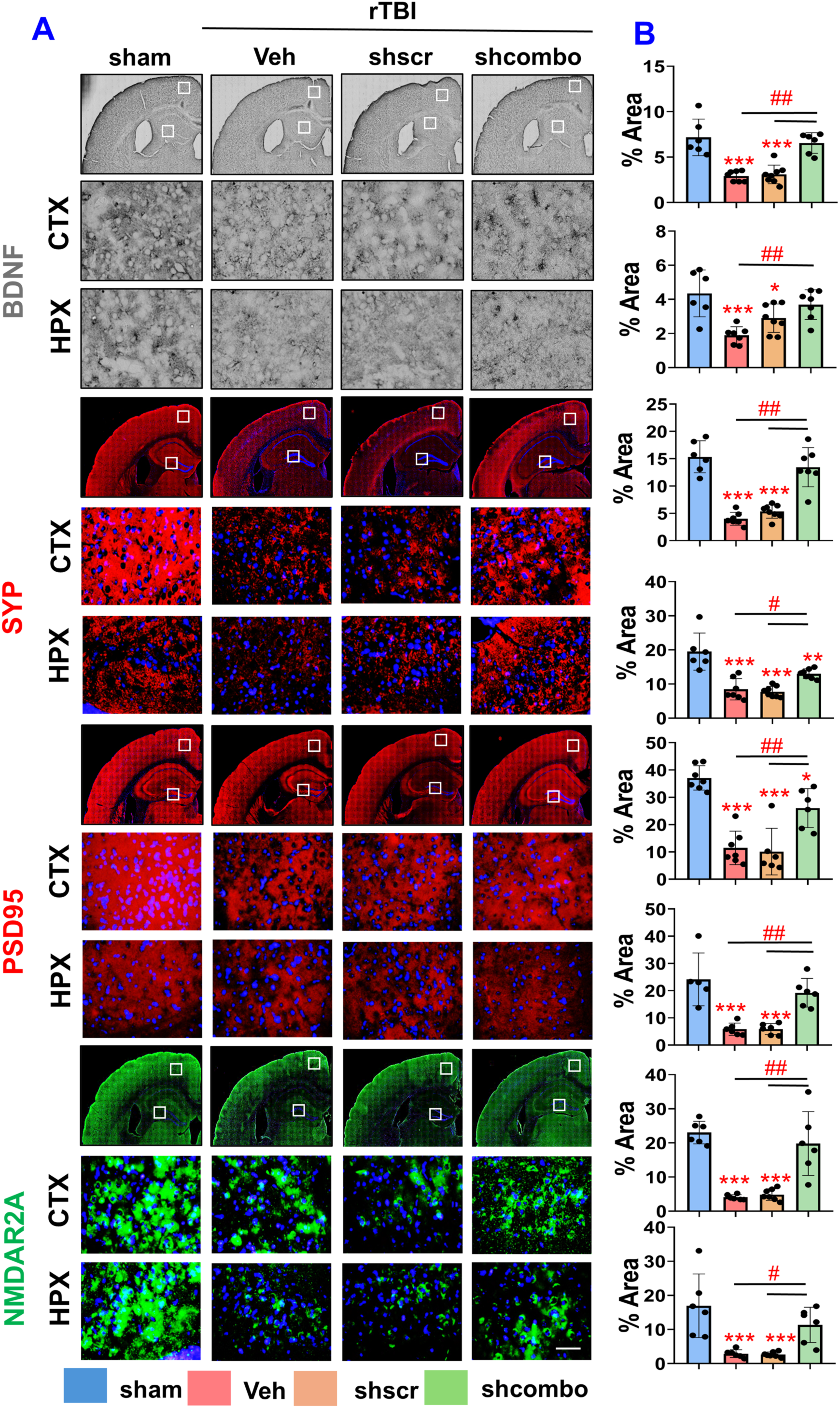
Downregulation of the CCL20–CCR6 axis alleviates rTBI⍰induced neuroinflammation and Protects Synaptic Integrity. **(A)** Brightfield and fluorescent images showing the expression of BDNF, Synaptophysin/SYP, PSD95, and NMDAR2 in the cortex (CTX) and hippocampus (HPX) counterstained with DAPI. **(B)** Quantification of BDNF, SYP, PSD95, and NMDAR2 expression (% area) in the CTX and HPX of rTBI mouse brains at 30 dpi. Scale bar: 50 µm (inset: 10 µm). n□ =□ 6/group. Data are presented as mean □±□ SD. Statistical analysis: one⍰way ANOVA with Holm–Sidak test; *p□ <□ 0.05, **p □<□ 0.01, ***p □<□ 0.001 vs. sham; #p □< □0.05, ##p□ < □0.01 vs. treated groups.

### CCL20 signaling drives microglial activation and serves as a therapeutic target for complement⍰mediated synaptic damage after rTBI

To determine whether CCL20 directly contributes to microglial activation and synaptic damage in vivo, recombinant CCL20 was administered to mice. Immunofluorescence staining revealed a significant increase in activated microglia in both the cortex and hippocampus of CCL20⍰injected mice, comparable to LPS⍰treated controls. Notably, these mice also exhibited sparse and fragmented synaptophysin⍰positive puncta at 14 days post⍰injection. In contrast, mice injected with mouse serum albumin (MSA) displayed uniform synaptophysin expression and minimal microglial activation, similar to sham controls (Fig 3A, B, S8). Further, IBA1/C1q double staining showed a substantial population of IBA1⍰positive microglia co⍰expressing C1q in the cortex and hippocampus of CCL20⍰ and LPS⍰treated mice, whereas MSA⍰injected and sham mice showed minimal C1q labeling. Together, these findings demonstrate that CCL20 is sufficient to drive microglial activation and complement dysregulation, leading to synaptic loss (Fig 3A, B, S8).

To further evaluate whether neutralizing CCL20 could rescue rTBI⍰induced pathology, mice were treated with an anti-CCL20 antibody. Immunostaining confirmed that CCL20 expression in the cortex and hippocampus was significantly reduced in anti-CCL20-treated animals compared with vehicle⍰ and isotype⍰treated rTBI mice, which showed robust CCL20 upregulation relative to sham controls (Fig 3C, D, S9). To assess whether CCL20 neutralization protects against microglia⍰mediated synaptic damage, we examined the co-expression of IBA1 and SYP. Consistent with previous findings, vehicle⍰ and isotype⍰treated rTBI mice exhibited marked reductions in SYP puncta, accompanied by increased clustering of IBA1+ microglia within the injured cortex and hippocampus. In contrast, anti⍰CCL20 treatment preserved SYP puncta and significantly reduced microglial density, resulting in an increased SYP area fraction and decreased IBA1 immunoreactivity (Fig 3C, D, S9). These results indicate that CCL20 neutralization preserves synaptic integrity by attenuating microglial activation. To determine whether CCL20 neutralization also modulates complement-mediated synaptic injury, IBA1/C1q co⍰immunostaining was performed. Vehicle⍰ and isotype⍰treated rTBI mice displayed extensive co⍰expression of IBA1 and C1q, evident as dense yellow puncta, consistent with complement⍰mediated synaptic loss. Anti⍰CCL20 treatment markedly reduced both IBA1+ microglial c density and C1q deposition, confirming that CCL20 downregulation mitigates complement activation and protects synaptic architecture in both the cortex and hippocampus (Fig 3C, D, S9).

### Treatment with shCombo⍰DPX mitigates glial activation and attenuates synaptic damage in rTBI mice

Building on our earlier findings demonstrating reduced neurodegeneration and glial activation at 7 DPI with shCombo⍰DPX treatment, we extended our analysis to 30 dpi in both the primary (cortex) and the secondary (hippocampus) injury site [29] (Fig 4A). Immunostaining revealed that vehicle- or shScr DPX⍰treated rTBI mice exhibited markedly elevated CCL20, CCR6, GFAP, and IBA1 expression in both regions, indicating persistent chemokine signaling and sustained glial activation long after repeated injury. In contrast, shCombo⍰DPX treatment substantially reduced CCL20 and CCR6 levels, accompanied by decreases in GFAP and IBA1 expression in both regions (Fig. 4B–E), demonstrating broad suppression of chronic neuroinflammation. Notably, microglia and astrocytes in shCombo⍰treated animals displayed a more ramified, resting⍰like morphology, in stark contrast to the hypertrophic, activated phenotypes observed in vehicle-treated rTBI mice. Morphometric analysis was performed to assess microglial phenotypic changes following shCombo DPX treatment. Vehicle- or isotype-treated rTBI mice showed a significant increase in fractal dimension, indicating chronic microglial activation. In contrast, shCombo DPX treatment significantly reduced fractal dimension, restoring a more ramified morphology consistent with a homeostatic microglial state. These structural changes were accompanied by reduced microglial clustering and improved synaptic integrity, demonstrating that shCombo DPX attenuates microglial hyperactivation and mitigates rTBI⍰induced synaptic loss (Fig. S10 A, B). Consistent with these findings, IBA1–CD68 double immunostaining revealed an increased population of IBA1-positive microglia co⍰expressing CD68 in vehicle- or isotype-treated rTBI mice, reflecting a persistently activated phenotype. In contrast, shCombo DPX treatment significantly reduced CD68 expression within IBA1+ microglia, showing diminished microglial activation (Fig. S10 C, D).

### Downregulation of the CCL20–CCR6 axis alleviates rTBI⍰induced neuroinflammation and Protects Synaptic Integrity in rTBI mice

BDNF, a key neurotrophic factor regulating neurogenesis and synaptogenesis, was markedly reduced in the cortex and hippocampus of rTBI mice treated with vehicle or shScr-DPX compared with sham controls (Fig. 5A, B). shCombo⍰DPX treatment significantly increased BDNF expression relative to both rTBI and shScr-DPX groups, indicating that suppressing CCL20–CCR6 signaling helps restore BDNF levels after injury. Presynaptic (synaptophysin, SYP) and postsynaptic (PSD95) markers were then assessed to determine whether attenuating CCL20–CCR6 protects against synaptic integrity. SYP puncta were significantly reduced in vehicle⍰ and shScr⍰DPX treated rTBI mice, whereas shCombo⍰DPX preserved SYP density in both the cortex and hippocampus (Fig. 5A, B). A similar pattern was observed for PSD95. Vehicle⍰ and shScr⍰DPX treated rTBI mice showed substantial loss of PSD95-positive puncta in both regions, while shCombo⍰DPX treatment preserved postsynaptic labeling, approaching sham levels (Fig. 5A, B). To further assess excitatory synapse integrity, NMDAR2A expression was assessed. Vehicle⍰ and shScr-DPX⍰treated rTBI mice showed a pronounced reduction in NMDAR2⍰positive puncta, whereas shCombo⍰DPX treatment significantly rescued NMDAR2A expression (Fig. 5A, B).

### Downregulation of the CCL20–CCR6 axis suppresses complement activation and synaptic pruning in rTBI mice

To assess whether CCL20 contributes to microglial activation, we performed IBA1/CCL20 double⍰immunofluorescence staining. A substantial population of activated microglia expressed CCL20 in both the cortex and hippocampus of vehicle⍰ and shScr⍰DPX-treated rTBI mice, whereas sham mice showed minimal signal. In contrast, shCombo⍰treated mice exhibited a marked reduction in CCL20 expression, accompanied by fewer IBA1⍰positive microglia in both regions (Fig. 6A, B, S11-12). To determine whether CCL20 localizes to presynaptic terminals, we performed SYP/CCL20 co⍰staining. CCL20 immunoreactivity was detected within synaptophysin⍰positive boutons, confirming its presence in neuronal compartments. Regions with high CCL20 expression consistently overlapped with areas showing reduced synaptophysin intensity, indicating that CCL20⍰rich microenvironments correspond to zones of synaptic vulnerability (Fig. 6A, B, S11-S12). This spatial relationship was pronounced in vehicle- and shScr⍰DPX-treated rTBI mice but greatly diminished in shCombo⍰treated mice, which showed increased synaptophysin signal and reduced CCL20 expression. Synaptophysin staining further revealed a pronounced loss of presynaptic integrity at 30 dpi. Vehicle⍰ and shScr-DPX⍰treated rTBI mice exhibited sparse, fragmented synaptophysin⍰positive puncta in both brain regions, replacing the dense, uniform bouton distribution observed in sham animals. These regions of persistent synaptic depletion contained significantly elevated numbers of activated microglia, particularly at the injury site (Fig. 6A, B, third panel). Treatment with shCombo⍰DPX markedly reduced this fragmented synaptic phenotype, resulting in a higher synaptophysin⍰positive area fraction and fewer activated microglia relative to vehicle⍰ or shScr-DPX-treated mice. To examine whether CCL20 downregulation influences complement activation, we performed IBA1/C1q co⍰staining. Vehicle⍰ and shScr⍰DPX-treated rTBI mice showed a substantial population of IBA1⍰positive microglia co⍰expressing C1q in both the cortex and hippocampus. In contrast, shCombo-DPX-treated mice exhibited a significant reduction in C1q-positive puncta, indicating decreased complement cascade activation (Fig. 6A, B, S11, S12).

**Figure 6.**
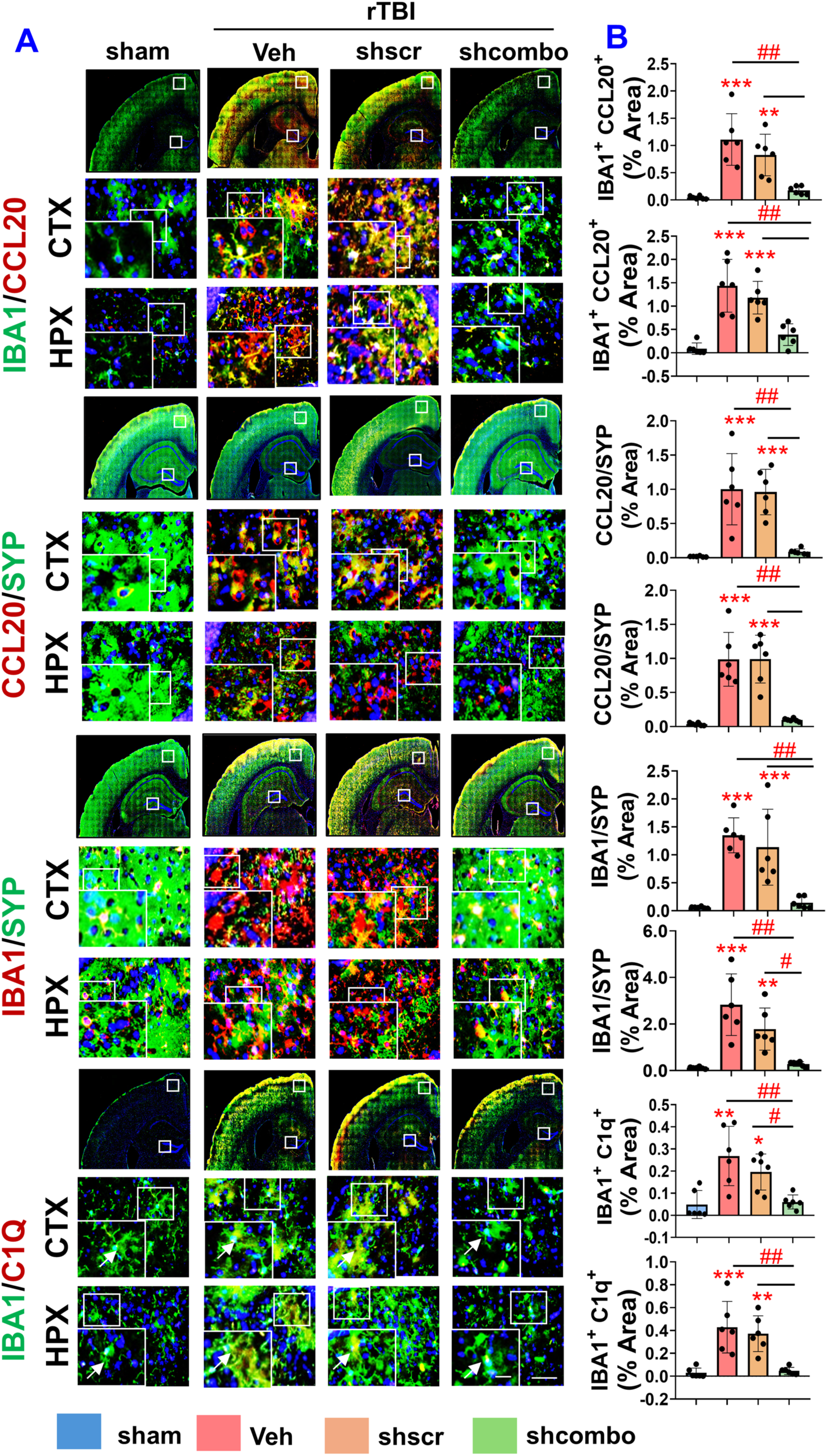
Downregulation of the CCL20–CCR6 axis suppresses complement activation and synaptic pruning in rTBI mice. **(A)** Representative immunofluorescence images showing double immunostaining of IBA1 (Green) / CCL20 (Red); SYP (Green) / CCL20 (Red); SYP (Green) / IBA1 (Red); IBA1 (Green) and C1Q (Red), counterstained with DAPI in the cortex (CTX) and hippocampus (HPX) at 30 DPI. **(B)** Histogram showing the quantification of the ratio of mean fluorescence intensity of IBA1/ CCL20, CCL20/ SYP, IBA1/ SYP, and IBA1+ C1q+ cells. (% Area). Scale bar: 50 µm (inset: 10 µm). Data are presented as mean ± SD. *p < 0.05, **p<0.01, ***p<0.001 vs. sham; ^#^ p < 0.05, ^##^p < 0.01, compared to treated groups; one⍰way ANOVA with Holm–Sidak test, n = 6/group.

Circulating levels of GFAP and C5a were measured in serum using ELISA. Serum GFAP concentrations were significantly elevated in rTBI mice treated with vehicle or shScr⍰DPX compared with sham controls. In contrast, rTBI mice treated with shCombo⍰DPX showed a marked reduction in circulating GFAP, indicating that dual CCL20/CCR6 silencing results in recovery and reduced glial reactivity (Fig. S13 A). Furthermore, C5a levels were significantly increased in rTBI mice receiving vehicle or shScr⍰DPX, consistent with complement activation after repeated injury. Treatment with shCombo⍰DPX significantly reduced serum C5a at 30 days post⍰injury, demonstrating that dual CCL20/CCR6 silencing effectively mitigates complement dysregulation associated with chronic post⍰traumatic neuroinflammation (Fig. S13 B).

### Downregulation of CCL20–CCR6 mitigates cognitive decline and anxiety-like behavior associated with rTBI-induced neurological deficits

To assess functional impairments caused by rTBI, we performed a battery of behavioral tests, including the OFT for locomotor activity and anxiety-like behavior, the NORT for recognition memory, and the Elevated Body Swing Test (EBST) for asymmetric motor coordination. All behavioral assessments were recorded and analyzed using ANY-maze video⍰tracking software. Baseline OFT measurements collected prior to rTBI showed no significant differences among groups in center-zone entries or total distance traveled (Fig. S14 A). However, at 30 dpi, vehicle- or shScr⍰DPX–treated rTBI mice displayed a marked reduction in center-zone entries and total distance traveled relative to sham controls, indicating heightened anxiety-like behavior. In contrast, shCombo-DPX-treated rTBI mice exhibited significantly increased center-zone exploration, indicating attenuation of rTBI⍰induced anxiety-like behavior (Fig. 7 A, B, S14 B). In the NORT, rTBI mice spent less time exploring objects compared to sham animals. Although total exploration time during the 2⍰h post⍰training session was comparable across groups, deficits emerged during the 24⍰h retention test. Vehicle- or shScr⍰DPX–treated rTBI mice showed a significantly reduced discrimination index (DI), whereas shCombo⍰DPX treatment restored DI values, indicating improved recognition memory and enhanced exploratory behavior. These findings demonstrate that downregulation of the CCL20–CCR6 axis ameliorates rTBI⍰induced cognitive impairments (Fig. 7 C, D). Motor coordination was evaluated using the EBST. Vehicle- or shScr⍰DPX–treated rTBI mice exhibited pronounced motor asymmetry, characterized by reduced lateral swings and prolonged immobility in the center. In contrast, shCombo⍰DPX–treated mice showed significantly increased swing activity, indicating improved motor function after rTBI (Fig. 7E).

**Figure 7.**
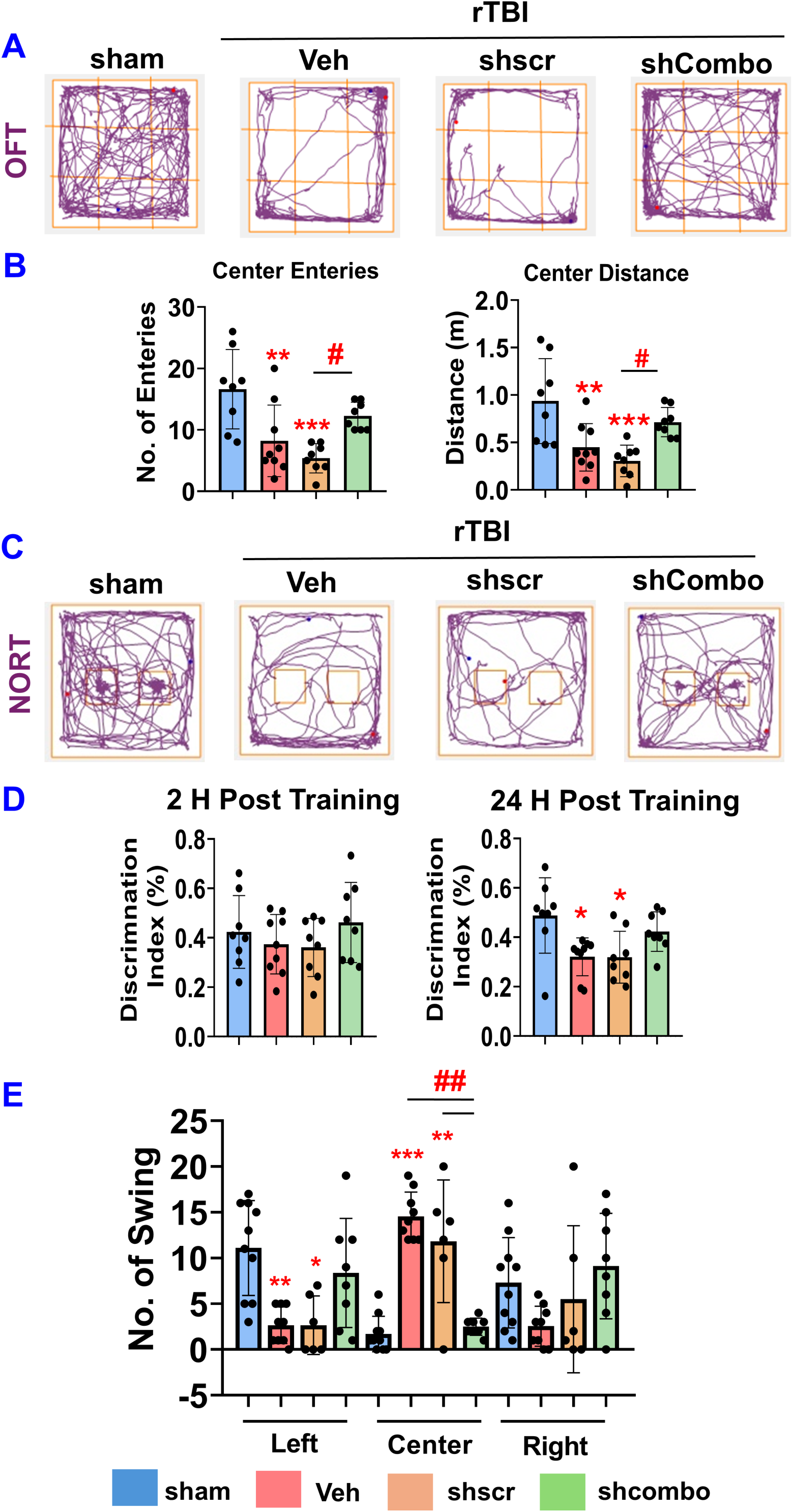
Downregulation of CCL20–CCR6 mitigates cognitive decline and anxiety-like behavior associated with rTBI-induced neurological deficits. **(A)** Representative track plots showing the mouse movement in the open field arena. **(B)** Histogram showing behavioral changes in the Open Field test (number of entries and distance traveled in the center zone) at 30DPI. **(C)** Representative track plots showing the mouse exposure to the familiar and novel objects. **(D)** Histogram depicting the discrimination index determined by the Novel Object Recognition test at 2 h and 24 h post-training. **(E)** Histogram representing asymmetric swing in the Elevated Body Swing Test (EBST) post-injury and treatment. Data are presented as mean ± SD. *p < 0.05, **p<0.01, ***p<0.001 vs. sham; ^#^ p < 0.05, ^##^p < 0.01, compared to treated groups; one-way ANOVA with Holm–Sidak test, n: sham=8; veh=9; shscr=8; shcombo=9 per group.

### Downregulation of CCL20 in IMG microglial cells attenuates indirect LPS-induced damage in HT22 neuronal cells

To investigate how microglia activation and secreted inflammatory factors influence neuronal health and synaptic integrity, we used a neuronal–microglial trans-well co-culture system (Fig. 8A). LPS⍰stimulated IMG cells showed a robust increase in fluorescence intensity, indicating strong upregulation of IBA1 and CCL20 following treatment with 500 ng/mL LPS. In contrast, IMG cells transfected with siCcl20 exhibited significantly reduced CCL20 and IBA1 expression, confirming effective knockdown and attenuation of microglial activation (Fig. 8A, B). HT22 neurons indirectly exposed to the secretomes of LPS⍰activated IMG cells for 48 hours displayed pronounced reductions in synaptic markers, including synaptophysin, demonstrating synaptic vulnerability driven by inflammatory microglial signaling. Notably, neurons co-cultured with CCL20⍰silenced IMG cells preserved higher levels of synaptophysin compared with neurons exposed to vehicle⍰ or siScr⍰treated IMG cells (Fig. 8C, D). Consistent with these findings, LPS stimulation increased CCL20 expression in vehicle⍰ and siScr⍰treated IMG cells, whereas CCL20 knockdown significantly reduced CCL20 levels in both IMG and HT22 cells. CCL20 silencing also prevented the LPS-induced reduction of the growth⍰associated protein GAP⍰43 in HT22 neurons. Furthermore, LPS⍰stimulated IMG cells induced caspase⍰3 activation and downregulated BDNF and PSD95 in vehicle⍰ or siScr⍰treated samples (Fig. 8E, F). In contrast, CCL20 knockdown reduced caspase⍰3 activation and significantly restored BDNF and PSD95 levels, highlighting the neuroprotective effect of reducing CCL20 in this co-culture model.

**Figure 8.**
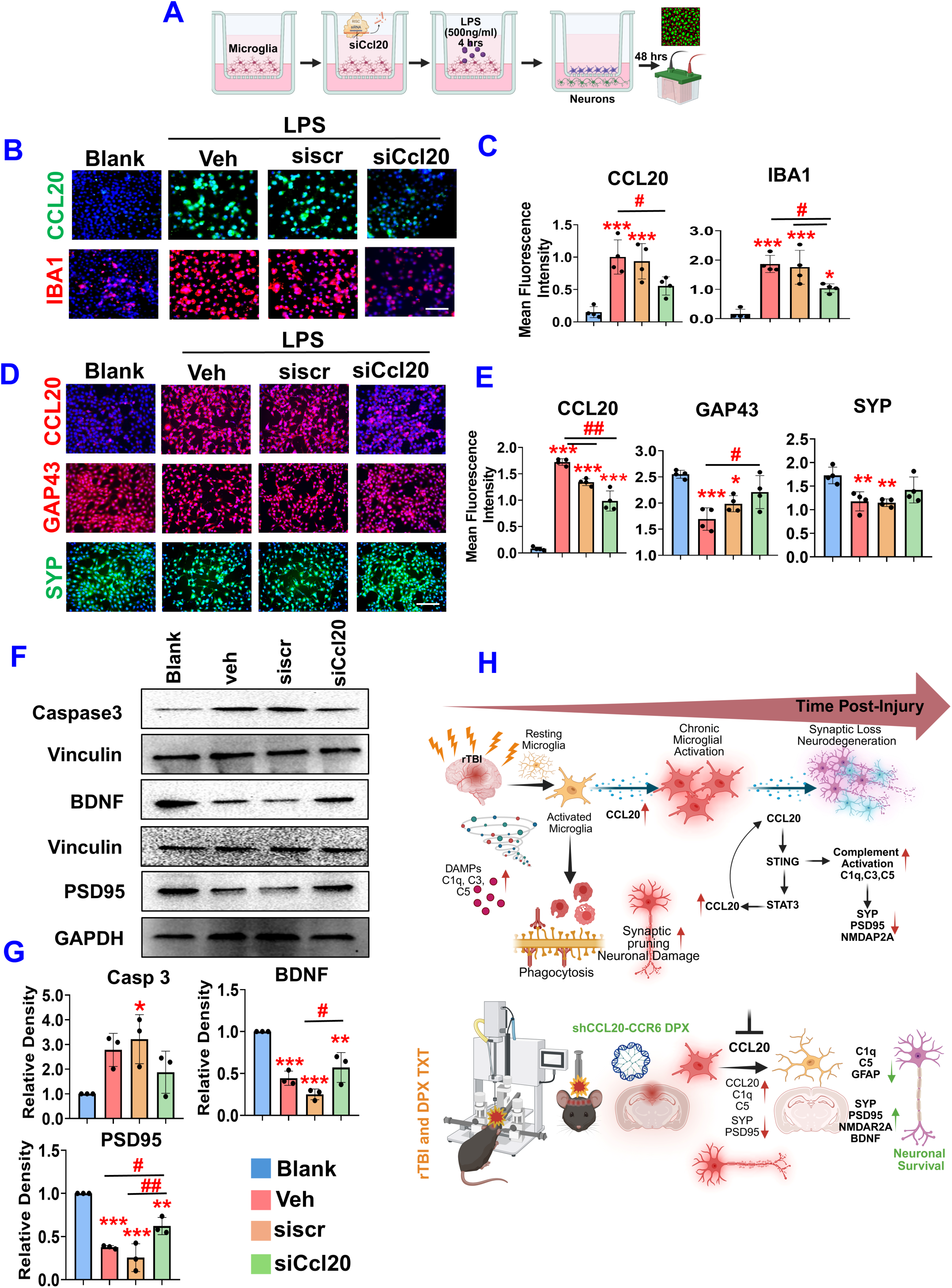
Downregulation of CCL20 in IMG microglial cells attenuates indirect LPS-induced damage in HT22 neuronal cells. **(A)** Schematic of the transwell co-culture experimental setup and timeline **(B)** Immunofluorescence images showing CCL20 and IBA1 staining in IMG cells stimulated with LPS (500 ng/ml) for 4 h, followed by culture in fresh media for 48 h. **(C)** Quantification of CCL20 and IBA1 immunofluorescence using ImageJ, expressed as mean fluorescence intensity. **(D)** Representative Immunofluorescence images of CCL20, GAP43, and SYP staining in HT22 cells exposed to secretome from LPS (500 ng/ml)-challenged IMG cells for 48h. **(E)** Quantification of CCL20, GAP43, and SYP immunofluorescence using ImageJ, expressed as mean fluorescence intensity, n = 4/group. **(F)** Representative immunoblots of synaptic plasticity (PSD95, BDNF) and apoptotic protein (Caspase-3). **(G)** Densitometric analysis of synaptic (PSD95, BDNF) and apoptotic protein (Caspase-3) expression normalized to vinculin or GAPDH, n=3/ group. Scale bar: 50 µm (inset: 10 µm). Data are presented as mean ± SD. *p < 0.05, **p<0.01, ***p<0.001 vs blank; ^#^ p < 0.05, ^##^p < 0.01, compared to treated groups; one⍰way ANOVA with Holm–Sidak test. **(H)**. Schematic representation illustrates the CCL20-STING-STAT3 signaling pathway hypothesis.

To directly assess the impact of CCL20 on synaptic integrity and neuronal health, HT22 cells were stimulated with recombinant CCL20 protein. CCL20 exposure also significantly decreased GAP43 immunoreactivity, indicating impaired neuronal growth (Fig S15 A, B). Consistently, protein levels of PSD95 and BDNF were markedly reduced at 48 hours post⍰treatment, confirming that CCL20 disrupts key markers of neuronal plasticity. In parallel, expression of the apoptotic marker caspase⍰3 was significantly elevated at 48 hours post-stimulation, demonstrating that CCL20 induces neuronal cell death (Fig. S15 C, D). Synaptophysin immunofluorescence revealed a significant reduction in dendritic length in CCL20⍰treated cells, with the most pronounced effect at 48 hours (Fig. S15 E, F).

## DISCUSSION

Repetitive TBI induces a chronic, self⍰perpetuating neuroinflammatory state in which unresolved immune activation drives progressive neurodegeneration and cognitive decline. The controlled cortical impact (CCI) model is widely used to investigate these pathological processes because it recapitulates key features of rTBI, including diffuse axonal injury, sustained microgliosis, astrogliosis, and behavioral impairments. These neuropathological and functional alterations closely resemble those observed in military veterans, athletes, and other individuals exposed to repeated head trauma. A central finding of this study is that the CCL20–CCR6 axis sustains chronic neuroinflammation by promoting microglia⍰mediated, complement⍰dependent synaptic degeneration. This represents the first demonstration that CCL20 signaling regulates complement⍰driven synaptic loss in the chronically injured brain at 30 dpi, thereby establishing a mechanistic link between chemokine activity and long⍰term synaptic vulnerability.

CCL20 is increasingly recognized as a clinically relevant inflammatory chemokine across a range of neurological disorders, including infectious conditions such as pneumococcal meningitis [36], and HIV⍰associated neuroinflammation [37], autoimmune and chronic inflammatory disorders such as multiple sclerosis [27], psoriasis, and rheumatoid arthritis [37], as well as neurodegenerative and ischemic disorders including AD, spinal cord injury, and stroke [24, 27, 36, 38, 39]. Consistent with this growing body of literature, prior work from our group demonstrated that CCR6⍰deficient mice or pharmacologic inhibition of CCL20 signaling via anti-CCL20 antibodies or PPARγ agonists such as pioglitazone and rosiglitazone significantly reduced neurodegeneration and gliosis while improving behavioral outcomes following rTBI [28, 29, 40–43].

Proteomic profiling at 30 dpi revealed dysregulation of pathways related to inflammation, complement activation, synaptic function, and stress responses, correlating with persistent microglial activation and synaptic injury. These sustained molecular alterations underscore the central role of chronic microglial activation and synaptic damage as hallmarks of rTBI [16–18]. Immunohistochemical analysis confirmed that markers of microglial activation, synapse-associated proteins, and IBA1–C1q colocalization persisted through 120 dpi, though at reduced levels compared to 30 dpi, indicating that 30 dpi represents a robust snapshot of secondary TBI pathology. These findings align with reports that primed microglia maintain prolonged cytokine, chemokine, ROS, and complement production [7, 44], contributing to maladaptive synaptic pruning and diffuse axonal injury [45, 46]. Chronic microgliosis has been consistently observed across animal models and human rTBI brains, [4, 47], where microglia remain activated for months to years, accompanied by systemic inflammation and long-lasting cognitive deficits.[48–51].

Mechanistically, complement deposition localized to activated microglia, consistent with trauma induced DAMP release, BBB disruption, and classical complement cascade activation [17, 52]. C1q binding to stressed synapses initiates C3 cleavage and opsonization, marking synapses for elimination via microglial complement receptors [19, 20, 53, 54]. Sustained cytokine signaling and C5a–C5aR1 engagement perpetuate hyperphagocytic microglial states, linking rTBI-induced synaptopathy to neurodegenerative trajectories observed in AD, Genetic or pharmacologic disruption of complement receptors attenuates microglial activation and synaptic tagging, underscoring their role as active regulators of the neuroimmune environment [21, 55] [56, 57]. C3aR-deficient mice display reduced astrogliosis and improved neuronal survival following traumatic or toxic insults [58], while C5aR1deficiency diminishes NF κB activation, leukocyte recruitment, and chronic microgliosis [59]. Collectively, these studies establish complement receptors as active regulators of the neuroimmune environment rather than passive downstream effectors and demonstrate that inhibiting complement signaling shifts the microglial response toward a less damaging phenotype [60].

Our findings demonstrate that shCCL20–CCR6 DPX therapy provides sustained suppression of microglial activation and protection against synaptic vulnerability, thereby establishing a clinically relevant therapeutic window for intervention. Consistent with previous reports, free shRNA exhibited limited stability and was rapidly degraded, whereas dendrimer-complexed shRNA achieved efficient and durable gene silencing [61]. Importantly, gene expression profiling revealed no significant changes in innate immune or inflammatory responses following DPX treatment up to 30 dpi, supporting immunological safety profile (Fig. S16, 17). This favorable safety profile was further corroborated by normal liver enzyme levels, complete blood counts, and histopathological analyses (Table S3-4). Collectively, these findings position DPX dendrimer therapy as a safe and effective nanoplatform with strong translational potential for mitigating neuroinflammation, and preserving synaptic vulnerability, and improving outcomes following TBI.

Silencing CCL20 disrupted CCR6-mediated recruitment of proinflammatory immune cells and attenuated microglial activation, thereby reducing complement deposition at stressed synapses. This suppression dampened C1q–C3 signaling, limiting opsonization and preventing excessive synaptic tagging and pruning. These findings are particularly relevant in the context of dynamic neuroimmune response to rTBI, where coordinated interactions among microglia, astrocytes, infiltrating leukocytes, and cytokine–chemokine networks shape acute, subacute, and chronic inflammatory phases, complicating effective immunomodulation[15, 62]. Collectively, our results identify CCL20 as an upstream regulator of complement activation and support the existence of a feed-forward CCL20–CCR6–STING–STAT3 signaling axis [63, 64] (Fig. 8 H) that sustains microglial reactivity, enhances synaptic tagging, and accelerates complement-dependent synaptic pruning. This framework provides a mechanistic link between chemokine-driven neuroinflammation and long-term synaptic vulnerability. Although further studies are required to delineate precise molecular interactions and neuronal contributions, our findings position CCL20–CCR6 signaling as a key driver of chronic synaptic degeneration and a promising therapeutic target in rTBI.

In conclusion, persistent CCL20 elevation and CCR6 activation amplify microglial complement signaling, creating a self⍰reinforcing inflammatory cascade that promotes progressive synaptic loss following rTBI. By identifying the CCL20-CCR6–complement axis as a critical mechanistic bridge between injury⍰induced inflammation and progressive neurodegeneration, our study advances current understanding of secondary injury. Moreover, the therapeutic benefits achieved through dendrimer-mediated CCL20 gene silencing highlight the translational potential targeted nanomedicine approaches for mitigating chronic pathology and preserving synaptic integrity after rTBI.

## MATERIALS AND METHODS

### Cell Culture, Transfection, and Stimulation Study

HT⍰22 mouse hippocampal neuronal cells (SCC129, Sigma⍰Aldrich, MO, USA) and IMG immortalized microglial cells (SCC134, Sigma⍰Aldrich, MO, USA) were maintained in Dulbecco’s Modified Eagle Medium (Gibco, NY, USA) supplemented with 10% fetal bovine serum (FBS) and 1% antibiotics (Hyclone, UT, USA) at 37°C in a 5% CO₂ incubator. To assess the direct effects of CCL20 on neuronal synaptic integrity, HT⍰22 cells were treated with 500 ng/mL recombinant CCL20 (760-M3, R&D Systems) for 24 or 48 hours and processed for immunostaining or western blot analysis. To examine microglial phenotypic changes and the impact of microglia⍰derived cytokines on neuronal synapses, HT⍰22 neurons (lower chamber) were co-cultured with IMG microglia (upper chamber) using 24⍰well and 6⍰well Transwell systems (Greiner, Bio-One, 657641, 662641). Smart⍰pool siRNAs targeting CCL20 (L⍰065734⍰01⍰0005) along with a scramble control (D⍰001810⍰10⍰05), were obtained from Horizon Discovery (Cambridge, USA). siRNA transfections were performed using the Mirus TransIT⍰X2 Dynamic Delivery System (MIR 6000, Madison, USA) following the manufacturer’s protocol. Twenty⍰four hours after transfection with either scramble siRNA (siScr) or siCcl20, IMG cells were stimulated with LPS (L2630, Sigma-Aldrich) (500 ng/mL) for 4 hours, washed with PBS, and the Transwell inserts were transferred onto HT⍰22 cultures for an additional 48⍰hour co-culture period. At the end of the experiment, HT⍰22 and IMG cells were fixed with 4% paraformaldehyde for immunostaining or lysed in RIPA buffer for western blotting.

### Animal Study, Induction of rTBI, and Therapeutic Regimen

All animal procedures followed NIH guidelines for the Care and Use of Laboratory Animals and were approved by the Institutional Animal Care and Use Committee at the University of South Florida. Male C57BL/6 mice (6–8 weeks old) were housed under a 12 h light/12 h dark cycle with food and water available ad libitum.

The initial studies were designed to examine the direct and chronic effects of CCL20. A small cohort of mice received intraperitoneal injections of recombinant CCL20 protein, while control cohorts received mouse serum albumin (negative control) or lipopolysaccharide (positive control). Treatments were administered at 0.5 mg/kg for two weeks (n = 3 per group).

For rTBI induction, mice were anesthetized with 2% isoflurane in oxygen and placed on a heating pad to maintain body temperature. The scalp over the impact site was shaved (∼1 cm), and the head was secured in a stereotaxic frame. An electromagnetic-controlled cortical impact (CCI) device was positioned at −0.8 mm anteroposterior from the bregma along the midline. Impacts were delivered using a 5-mm metal tip at 5 m/s, with a depth of 1.5 mm and a dwell time of 200 ms, covering approximately 1.7 mm to −3.3 mm relative to the bregma. Mice received five impacts, one every 48 hours. No skull fractures, contusions, or hemorrhages were observed, and no mortality occurred during rTBI induction or treatment. The experiment was repeated twice, with n = 6 per group for immunopathological analyses and n = 3 per group (sham and rTBI) for proteomic analyses.

For treatment, shRNA plasmid DNA (shscr DPX or shcombo DPX) was administered intranasally and intravenously twice weekly at 7 and 21 days post-injury. In another subset of rTBI-induced mice, 20 µg of mouse monoclonal anti-CCL20 antibody (MAB760, R&D Systems) or isotype control (MAB005, R&D Systems) was administered intraperitoneally at 1 mg/kg every other day for a total of five doses.

30 days after the final injury, animals were deeply anesthetized with Euthasol (150 µg/mL, i.p.). Blood was collected, and mice were perfused transcardially with 0.1 M phosphate buffer followed by 4% paraformaldehyde. Brains were harvested, post-fixed in 2% paraformaldehyde, and cryoprotected in 30% sucrose. Coronal sections (30 µm) were cut on a cryostat. Blood samples were allowed to clot at room temperature for 30 minutes, centrifuged at 2000 × g for 15 minutes, and serum was stored at −80 °C. Additional blood was collected in heparinized tubes for RNA isolation using the Takara NucleoSpin RNA Blood kit.

### Preparation and Evaluation of shDendriplexes

The preparation and characterization of DPX was performed as previously described by Mayilsamy et al. (2020). Briefly, dendrimers containing 10 µg of pDNA were prepared in a final volume of 50 µL in water, with 25 µL delivered into each nostril under isoflurane anesthesia. For intravenous administration, dendrimers containing 10 µg of pDNA were prepared in a final volume of 100 µL and injected via the tail vein. Particle size and zeta potential of the DPX formulation were measured using a Zetasizer prior to administration to confirm stability and reproducibility. The shRNA target sequences were designed and potential off⍰target effects were evaluated through bioinformatic analysis; plasmid details are provided in the supplemental information (Table S5). To assess biodistribution, fluorescently labeled DPX was administered and tracked following intravenous versus intranasal delivery. Systemic safety was evaluated through liver enzyme assays, complete blood counts, and histopathological analyses, while immune responses triggered by nanoparticle administration were examined to establish the overall safety profile of the formulation.

### Behavioral Study

#### Open Field Test

Anxiety⍰like behavior and locomotor activity were assessed using the open field test (OFT). Mice were introduced into a square open⍰field arena (90 × 90 × 40 cm) subdivided into nine equal zones consisting of corner, perimeter, and center areas. Animals were gently placed in the center of the arena and allowed to acclimate for 1 minute before testing. Each mouse was then tracked for 10 minutes using a video camera, and locomotor activity and zone⍰specific behavior were analyzed with Anymaze software. The arena was cleaned thoroughly with 75% ethanol between trials to eliminate olfactory cues.

#### Novel object recognition Test

Hippocampal⍰dependent recognition memory was assessed using the novel object recognition test (NORT). On the second day following exposure to the open⍰field arena, mice were placed in the arena containing two identical objects (A and A) and allowed to explore for 5 minutes. After a 5⍰minute break, this familiarization session was repeated twice for a total of three 5⍰minute exposures. 2 hours and 24 hours after the initial training session, mice were returned to the arena and exposed to one familiar object (A) and one novel object (B) for 10 minutes, with behavior recorded. After each set of mice, the objects and boxes were thoroughly cleaned with 70% ethanol to avoid olfactory cues. The recorded videos were analyzed using AnyMaze software. The discrimination index (DI) was calculated as the ratio of the novel object’s exploration time to the total exploration time. * Exploration time was defined by the animal sniffing or touching the object.

#### Elevated Body Swing Test

The elevated body swing test (EBST) was used to assess asymmetric motor behavior in mice. Each mouse was briefly placed in an empty cage and allowed to habituate for 1 minute. The test began by holding the mouse in a neutral position, with all four paws on the ground, then gently lifting it by the base of the tail to a height of approximately 1–2 inches. A swing was recorded when the mouse moved its body more than 10° to either the left or right. If the mouse remained immobile for more than 10 seconds, it was scored as maintaining a center position. A total of 20 swings were recorded for each animal. All sessions were videotaped and scored manually.

#### Isolation of Mouse Microglia

To determine whether microglia are the major cellular source of CCL20 following rTBI, sham and rTBI mice underwent microglial isolation. After euthanasia and transcardiac perfusion with ice⍰cold PBS, brains were removed and cortical/hippocampal regions dissected. Tissue punches (20–50 mg) were minced and enzymatically digested using the gentleMACS Octo Dissociator with manufacturer⍰specified enzyme mixes to generate single⍰cell suspensions. The homogenates were filtered through 70 µm strainers, and debris, myelin, and red blood cells were removed by gradient centrifugation and lysis. The resulting cell pellet was resuspended in PB buffer and incubated with CD11b⍰conjugated magnetic microbeads for 15 minutes at 4 □°C. Labeled cells were applied to MS columns in a magnetic separator, washed, and eluted to obtain purified CD11b⁺ microglia. Both the isolated microglial population and the remaining cell fraction were collected for RNA extraction and subsequent gene expression analysis.

#### RNA Isolation and Polymerase Chain Reaction

Total RNA was isolated using TRIzol (Life Technologies). To eliminate residual genomic DNA, RNA samples were treated with DNase I (Invitrogen, cat. no. 18068). One microgram of RNA was used for cDNA synthesis using the Maxima Enzyme 5× reaction mix (Thermo Fisher Scientific). Quantitative real⍰time PCR was performed on a CFX384 Touch™ Real⍰Time PCR Detection System (Bio⍰Rad). Each 5 µL reaction contained 1 µL of 5× qPCR master mix, 0.5 µL each of forward and reverse primers (Table S6), 1 µL of nuclease⍰free water, and 1 µL of cDNA. The cycling conditions were: 95°C for 3 minutes, followed by 45 cycles of 95°C for 10 seconds, 60°C for 1 minute, and 72°C for 15 seconds. All samples were run in triplicate across three independent experiments. Gene expression for each age group was normalized to its respective mock control.

#### Proteomic profiling

Cortex and hippocampus tissues were microdissected from sham and rTBI mice 30 days post injury. All samples were processed and analyzed at the Multi-Omics Data Facility within the USF Advanced Research Core for Mass Spectrometry. The tissue was physically homogenized, and a representative portion of each specimen was lysed and prepared using the iST sample preparation kit (PreOmics, GMBH). Briefly, tissue was probe sonicated in 100 µL iST LYSE buffer with protein extraction beads (Diagenode) then heated at 95 °C for 10 minutes. After extraction, protein concentrations were normalized to 50 µg in equal volumes digested in iST digest buffer. The resulting peptides were washed and purified using the iST cartridge. Peptides were analyzed by diaPASEF on a timsTOF Pro mass spectrometer coupled to a nanoElute 2 UHPLC system equipped with an Aurora Ultimate CSI reversed-phase C18 column (25 □cm × 75□ µm i.d., 1.7 □µm C18, IonOpticks, Fitzroy, Australia) heated at 50 °C. DIA-NN v. 2.3.0 was used for DIA data searching and quantification, and MS-DAP v. 1.2.2 was used for differential abundance analysis of the DIA-NN output. For DIA-NN, the --matrix-spec-q 0.01 command was utilized to achieve an additional run-specific <1% FDR filter at the protein level along with the standard global <1% protein and precursor FDR cutoff. A predicted library was generated from the Mus musculus UniProt database (downloaded June 2025, 63208 entries) with mass accuracy and MS1 accuracy both set to 15 ppm in the search parameters in addition to the remaining default settings. Differential abundance analysis was conducted in MS-DAP (v. 1.2.2) within RStudio [65] where the DIA-NN report file and FASTA file were specified and associated with sample metadata. The contrasts were set for differential abundance analysis using the algorithm MSqRob [66]. Minimum detection and quantification filters of at least 3 out of 4 replicates per group were used. Additionally, variance Stabilizing Normalization (VSN) and then Mode Between protein (MBprot) normalization steps were utilized prior to differential abundance analysis. The MSqRob p-value threshold was set at 0.01 and the log2 foldchange threshold was set to automatically infer through bootstrapping. Although q-values were calculated through Benjamini-Hochberg correction, the number of replicates (n=4) limited statistical power for proteomics-based discovery. Therefore, combined log2 foldchange and p-value thresholds were applied to enhance sensitivity while still accounting for FDR control.

Significantly differentially expressed (for both up- and downregulated) proteins (p≤0.05) were analyzed for functional enrichment and interactome relationships using Metascape and IPA [68]. Mouse protein IDs were converted into Entrez Gene IDs and analyzed as mouse based on both EggNOG and Homologene databases. A Circos plot was created to visualize protein overlap between cortex and hippo sample groups. Each arc on the plot represents a gene list, where each gene member of that list is assigned a spot on the arc. Dark orange color represents the genes that are shared by multiple lists and light orange color represents genes that are unique to that gene list. Purple lines link the same gene ID that is shared by multiple lists (a gene ID that appears in two lists will be mapped onto each arc once and the two positions are linked with a purple line). Blue lines link gene IDs that, although different, fall under the same ontology term. Enriched ontology clusters were visualized by heatmap. We first identified all statistically enriched terms (GO/KEGG terms, canonical pathways, hallmark gene sets, etc) using accumulative hypergeometric p-values with a p-value cutoff of 0.05 and enrichment factors were calculated and used for filtering. Background proteins for enrichment were set to the full list of proteins identified by mass spectrometry. Remaining significant terms were then hierarchically clustered into a tree based on Kappa-statistical similarities among their gene memberships. Then a 0.3 kappa score was applied as the threshold to cast the tree into term clusters. Terms with the lowest p-value within each cluster were selected as its representative term and displayed in a dendrogram. Protein-protein Interaction Enrichment was performed using the physical core database within Metascape. All protein-protein interactions among input genes were extracted from PPI data sources (OmniPath, InWeb_DB, BioGrid, STRING) and formed into a PPI network. GO enrichment analysis was applied to the network to extract biological meanings. The MCODE algorithm was applied to this network to identify neighborhoods where proteins are densely connected. Each MCODE network was assigned a unique color. GO enrichment analysis was applied to each MCODE network to extract biological meanings from the network component, where the top three lowest p-value terms were retained. Connections between Ccl20 and proteins within this network were plotted based on the STRING database (confidence score cutoff 0.5).

#### Western Blotting

Treated cells were washed with ice⍰cold PBS and gently scraped using a cell scraper. The cell suspension was transferred to microcentrifuge tubes and lysed in RIPA buffer (pH 7.4; Thermo Fisher Scientific, Waltham, MA, USA) supplemented with a protease and phosphatase inhibitor mix (Halt) Lysates were vortexed briefly and incubated on ice for 15 minutes, then clarified by centrifugation at 20,000 × g for 30 minutes at 4°C. The supernatant was collected, and protein concentration was measured using the Pierce Coomassie protein assay kit. Twenty micrograms of protein or molecular weight standards (Thermo Fisher Scientific, Waltham, MA, USA) were separated by SDS⍰polyacrylamide gel electrophoresis (Bio⍰Rad, Hercules, CA, USA) and transferred to a 0.2 µm nitrocellulose membrane (Bio⍰Rad, Hercules, CA, USA) at 80 V for 20 hours. Membranes were blocked for 1 hour at room temperature with 5% milk in TBS⍰T, or 5% bovine serum albumin for phosphoprotein detection. Blots were incubated with primary antibodies (Table S8) overnight at 4°C with gentle rocking, followed by incubation with the appropriate HRP⍰conjugated secondary antibodies for 2 hours at room temperature. After three washes in TBS⍰T (5 minutes each), signals were developed using SuperSignal™ West Pico PLUS chemiluminescent substrate (Thermo Fisher Scientific, Waltham, MA, USA) and imaged using the ChemiDoc XRS™ system (Bio⍰Rad, Hercules, CA, USA).

#### Enzyme-Linked Immunosorbent Assay (ELISA)

Blood was collected by standard cardiac puncture, and serum was separated by centrifugation at 2,000 × g for 10 minutes. The isolated serum was used to measure circulating GFAP using the Mouse Glial Fibrillary Acidic Protein (GFAP) ELISA Kit (RK09309) and C5a using the Mouse Complement C5a ELISA Kit (ab193718). All ELISAs were performed according to the manufacturer’s instructions.

#### Immunofluorescence Staining

IMG and HT22 cells were washed and fixed with 4% PFA for 10 minutes at room temperature. After three gentle PBS washes, cells were permeabilized with 0.2% Triton X⍰100 in PBS for 10 minutes. Cells were then washed, blocked with 5% bovine serum albumin (BSA) for one hour at room temperature, and incubated overnight at 4°C with primary antibodies diluted in 1% BSA prepared in PBST (0.1% Tween⍰20 in PBS). Following incubation, cells were washed and incubated with fluorescent⍰tagged secondary antibodies (Vector Laboratories, 1:1000, Supplementary Table S7) diluted in 1% BSA⍰PBST for 2 hours at room temperature, protected from light. Cells were washed again, counterstained with DAPI (Fisher Scientific, 1:1000) for 5 minutes at room temperature, washed, and imaged on an Olympus X71 microscope using appropriate filters.

Slide⍰mounted 30 µm brain sections were heated in antigen retrieval solution (1:100; Vector Laboratories, Burlingame, CA, USA) for 45 minutes at 90°C, cooled to room temperature, and washed with PBS. Sections were permeabilized and blocked for 1 hour in serum⍰blocking solution (10% host serum, 0.2% Triton X⍰100 in PBS). They were then incubated overnight at 4°C with primary antibodies (Table S7) prepared in 5% host serum and 0.1% Triton X⍰100 in PBS. After PBS washes, sections were incubated with fluorescently labeled secondary antibodies, washed again, air⍰dried, and mounted using DAPI⍰containing anti⍰fade mounting medium.

#### Immunoperoxidase Staining

For immunoperoxidase staining, sections underwent heat⍰mediated antigen retrieval followed by incubation in 3% hydrogen peroxide for 20 minutes to quench endogenous peroxidase activity. After blocking with a serum⍰blocking solution, sections were incubated overnight at 4°C with primary antibodies (Table S7). Following PBS washes, sections were incubated with biotinylated secondary antibodies for 2 hours at room temperature, then with avidin–biotin–peroxidase complex (ABC, 1:100; Vector Laboratories) for 1 hour. The signal was developed using a 3, 3′⍰diaminobenzidine (DAB) substrate (Vector Laboratories) for 5 minutes. Sections were washed, air⍰dried, and coverslipped with DPX mounting medium. Bright⍰field or fluorescence images were acquired using an Olympus X71 microscope with appropriate filters.

#### Image Analysis and Quantitation

All image quantification was performed using NIH ImageJ software. Immunohistochemical images were acquired using an Olympus IX71 microscope controlled by DP70 Manager Software (Olympus America, Melville, NY, USA). For each antibody, four coronal brain sections were analyzed, and six images per mouse were captured at 20× magnification. All images were collected using identical exposure and digital gain settings to minimize variability in the background signal. For DAB⍰based chromogenic IHC, sections were imaged using brightfield microscopy on the Olympus IX71 system under fixed exposure conditions. Fluorescently labeled sections were imaged at 20× magnification using a Keyence microscope (BZ-X710, Keyence America, Itasca, IL, USA), maintaining uniform exposure, minimizing background noise, and using the appropriate filter sets for DAPI (blue), Alexa Fluor 488 (green), and Alexa Fluor 594 (red).

Image analysis was performed using NIH ImageJ. Single⍰marker fluorescent images (Brain-derived neurotrophic factor (BDNF), N-methyl-D-aspartate receptor subunit 2A (NMDAR2A), Synaptophysin (SYP), Postsynaptic Density Protein (PSD95), CCL20) were converted to 8⍰bit grayscale, and a manual threshold (0–255) was applied to identify positively stained regions. Fluorescence intensity (integrated density) and percent area (% area) were quantified using the Analyze > Measure function.

For dual⍰marker analyses (MAP2/Iba1, SYP/Iba1, P2RY12/Iba1), the RGB channels were separated and analyzed individually to minimize background interference. Each channel was converted to 8⍰bit grayscale, thresholded (0–255), masked, and quantified for integrated density and % area. Microglia⍰associated neuronal or synaptic changes were assessed by calculating the ratio of Iba1⍰positive % area to the corresponding marker⍰positive % area. For CCL20/SYN staining, the ratio of CCL20⍰positive % area to SYN⍰positive % area was determined.

Co-expression for Iba1/C1q and Iba1/CCL20 was quantified by adjusting the threshold of overlapping signal (visualized as yellow in merged images) using the Adjust > Color Threshold tool. Brightness and contrast were optimized to reduce background noise, and % area of colocalized signal was measured using Analyze > Measure.

Images were converted to grayscale prior to analysis, and thresholds were adjusted uniformly across all samples to exclude background noise. Positive cells were manually counted for IBA1. For SYP, PSD95, NMDAR2, BDNF, C1Q, and GFAP staining, immunoreactivity was quantified as integrated density per unit area.

Morphological analysis of microglia was performed using fractal analysis. Photomicrographs were converted to grayscale, corrected for outliers, and thresholded to binary images. Individual microglia (five cells per photomicrograph) were randomly selected, converted to binary and outline images, and analyzed with the FracLac plugin in ImageJ using the box⍰counting method. Grid positions were set to six, and complementary parameters were selected. Final values were averaged and plotted. Synaptophysin-immunostained fluorescence images were acquired, and dendritic measurements were performed using ImageJ software. Individual neurons were selected, and dendritic processes were traced manually from the soma to the distal tip. The resulting values were averaged and expressed as mean dendritic length per neuron ± SEM. For each experimental group, 10 neurons were randomly selected from fields imaged at 20× magnification (n = 10 per group), yielding a total of approximately 100 cells analyzed across conditions.

To minimize the bias, all the analysis was performed in a blinded to the experimental conditions.

#### Statistical Analysis

All data are presented as mean ± SEM. Statistical significance was assessed using one⍰way or two⍰way ANOVA followed by the Holm–Sidak post hoc test for multiple comparisons unless otherwise specified. A p⍰value < 0.05 was considered statistically significant.

## DATA AVAILABILITY

Raw data and detailed analyses supporting the findings of this study are available from the corresponding author upon reasonable request. All other data generated or analyzed during this study are included in this published article (and its supplemental information).

## ACKNOWLEDGEMENTS

This work was supported by a Veterans Affairs Merit Review grant BX005757 awarded to Dr. Subhra Mohapatra, and by Research Career Scientist Awards to Dr. Subhra Mohapatra (IK6BX004212) and Dr. Shyam Mohapatra (IK6BX006032). Although this report is based in part on work supported by the Department of Veterans Affairs, Veterans Health Administration, Office of Research and Development, the contents do not represent the views of the Department of Veterans Affairs or the United States Government.

We would also like to acknowledge Jared Balkin and Premalata Pati for their valuable assistance in animal studies, behavioral scoring, tissue processing, and immunohistochemistry.

## AUTHOR CONTRIBUTIONS

SM, SSM, and AW designed the experiments. KM performed the experiments and wrote the manuscript. EM synthesized the nanoparticles, and PP assisted with tissue processing, immunostaining, and data analysis. SS, JG, and TW conducted the proteomics and analyzed the resulting datasets. RG performed the IPA analysis. SB and KT carried out the behavioral studies. SM, SSM, AW, EM, and SS reviewed the manuscript.

## DECLARATION OF INTERESTS

KM, EM, SSM and SM are inventors in a US patent (12, 589, 133 B2) covering the use of CCL20-targeting ShRNA

## Notes

### Competing Interest Statement

The authors have declared no competing interest.

