## Supplemental Information for "Dendrimer Delivered shRNA Targeting the CCL20–CCR6 Axis Suppresses Complement-Mediated Microglial Synaptic Pruning and Ameliorates Chronic Neuroinflammation After Repetitive Traumatic Brain Injury"

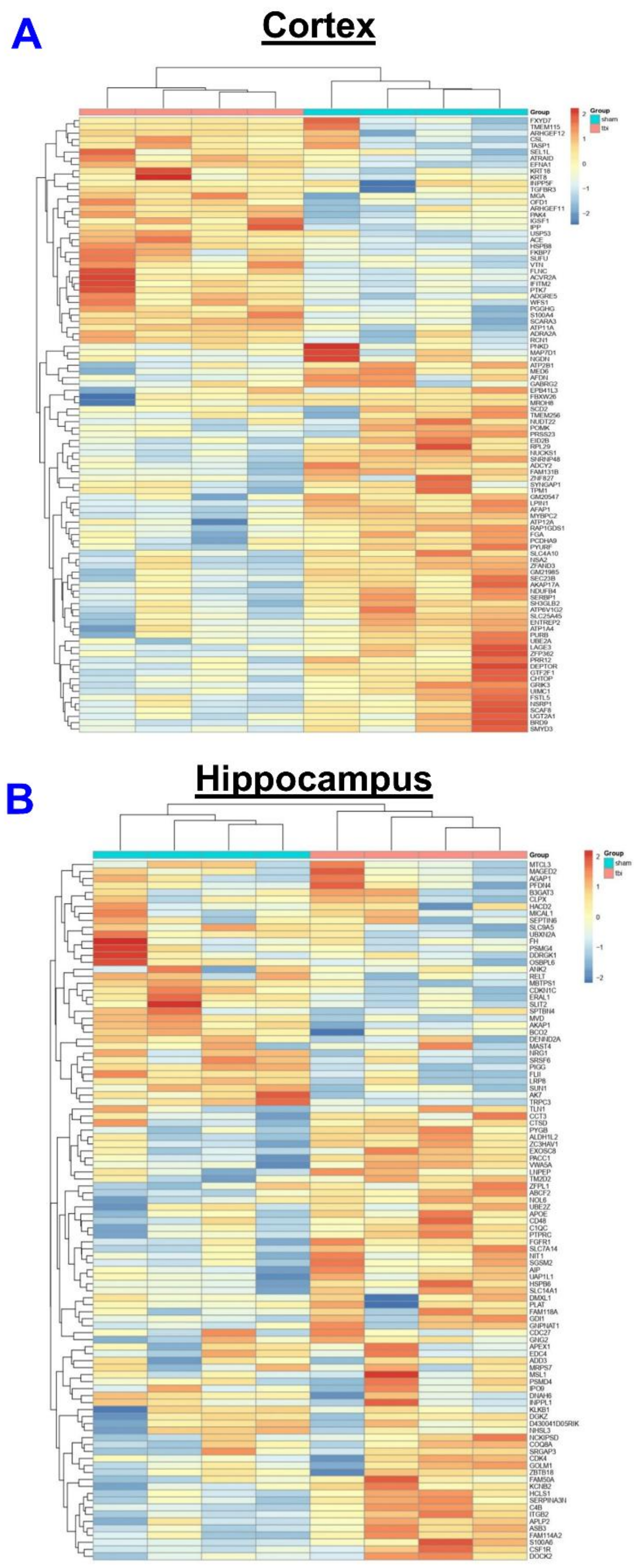

**Figure S1. Proteomic alterations in the cortex and hippocampus 30 days after rTBI**. Heat map representing differentially expressed proteins in the cortex (CTX) **(A)** and hippocampus (HPX) **(B)** of rTBI mice

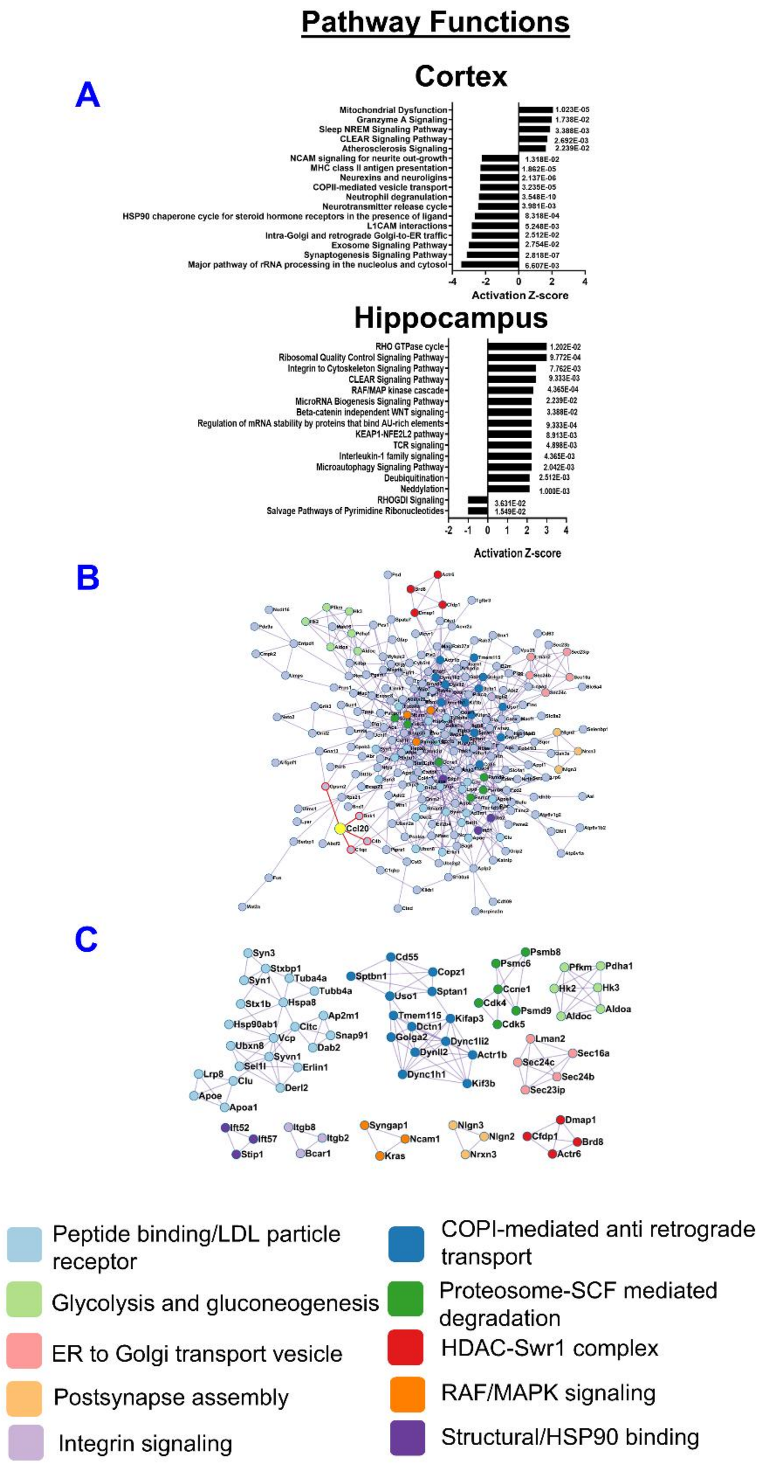

**Figure S2. Proteomic alterations in the cortex and hippocampus 30 days after rTBI.** Histograms summarizing pathway functions associated with categories enriched among differentially expressed proteins in the cortex (CTX) and hippocampus (HPX) (A) of rTBI mice (B) PPI network of the union of CTX and HPX DEPs. CCL20 was added to the network, and edges directly connected to CCL20 are highlighted in red. (C) Highly interconnected sub-regions identified by MCODE are color coded according to their top GO functional enrichment.

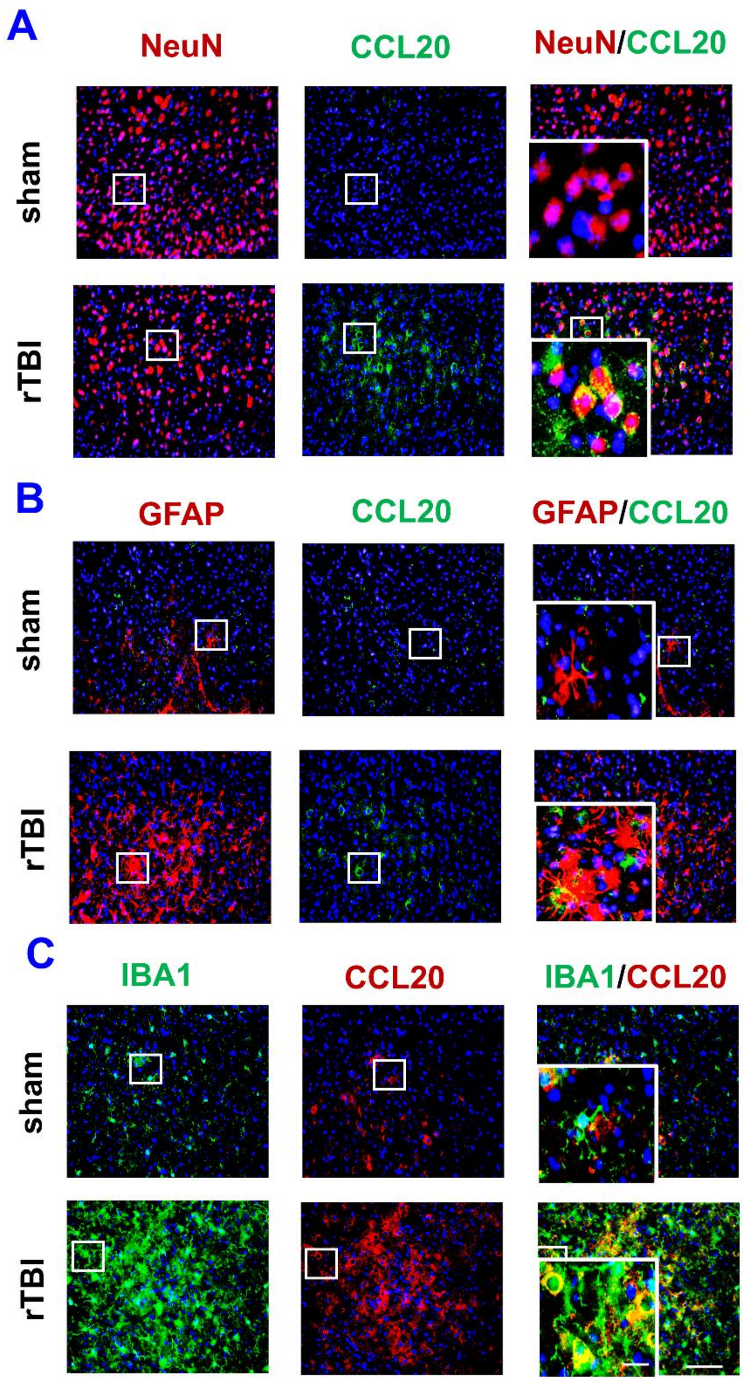

**Figure S3. Cell type-specific validation confirms microglia as the predominant CCL20 source after rTBI.** Immunostaining of CCL20 expression within neurons (NeuN+) (Red) **(A)**, astrocytes (GFAP+) (Red) **(B)**, and microglia (IBA1+) (Green) **(C)**, counterstained with DAPI in the cortical injury site at 30 dpi. Scale bar: 50 μm (inset: 10 μm).

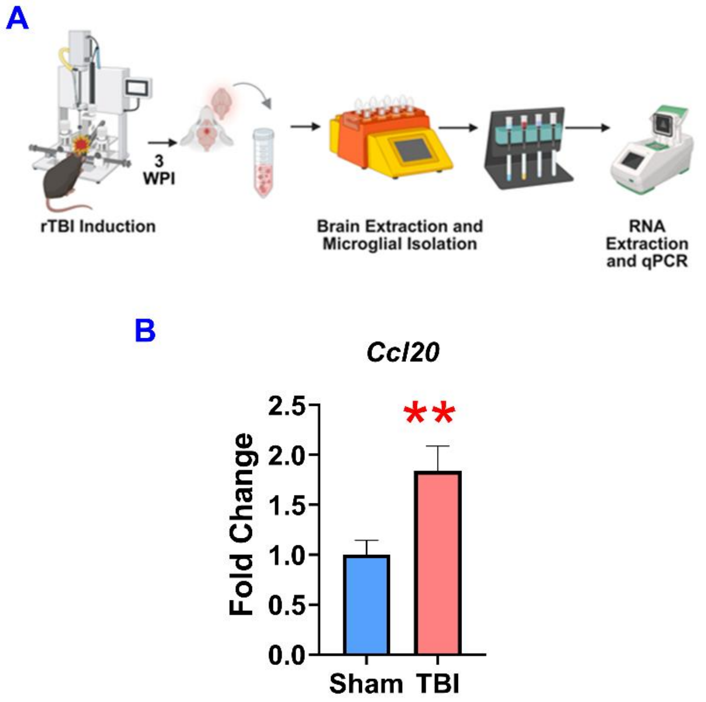

**Figure S4. Increased CCL20 expression in microglia isolated from rTBI mouse brains. (A)** Experimental design for isolating microglial cells from rTBI mouse brains. **(B)** qPCR analysis of Ccl20 expression in RNA isolated from microglia at 30 dpi ($n = 5$). Data are presented as mean ± SD. Statistical analysis: Student's test; *$p < 0.05$, **$p < 0.01$, ***$p < 0.001$ vs. sham

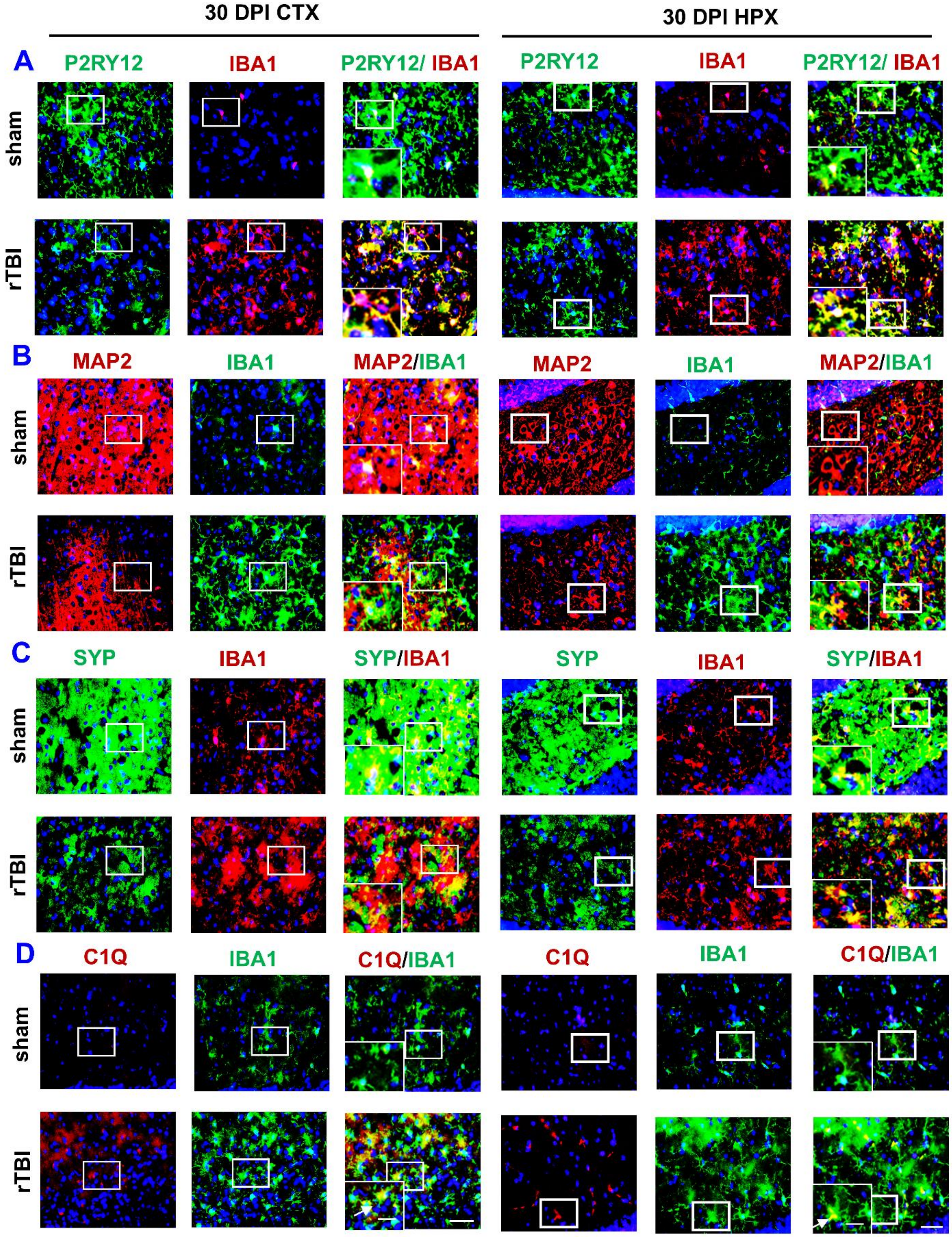

**Figure S5. rTBI induces chronic microglial activation and associated synaptic and neuronal damage at 30 dpi in rTBI mice.** Representative immunofluorescence images with individual channels showing double immunostaining of **(A)** P2RY12 (Green) / IBA1 (Red); **(B)** IBA1 (Green) / MAP2 (Red); **(C)** SYP (Green) / IBA1 (Red); **(D)** IBA1 (Green) and C1Q (Red), counterstained with DAPI in the cortex (CTX) and hippocampus (HPX) at 30 DPI

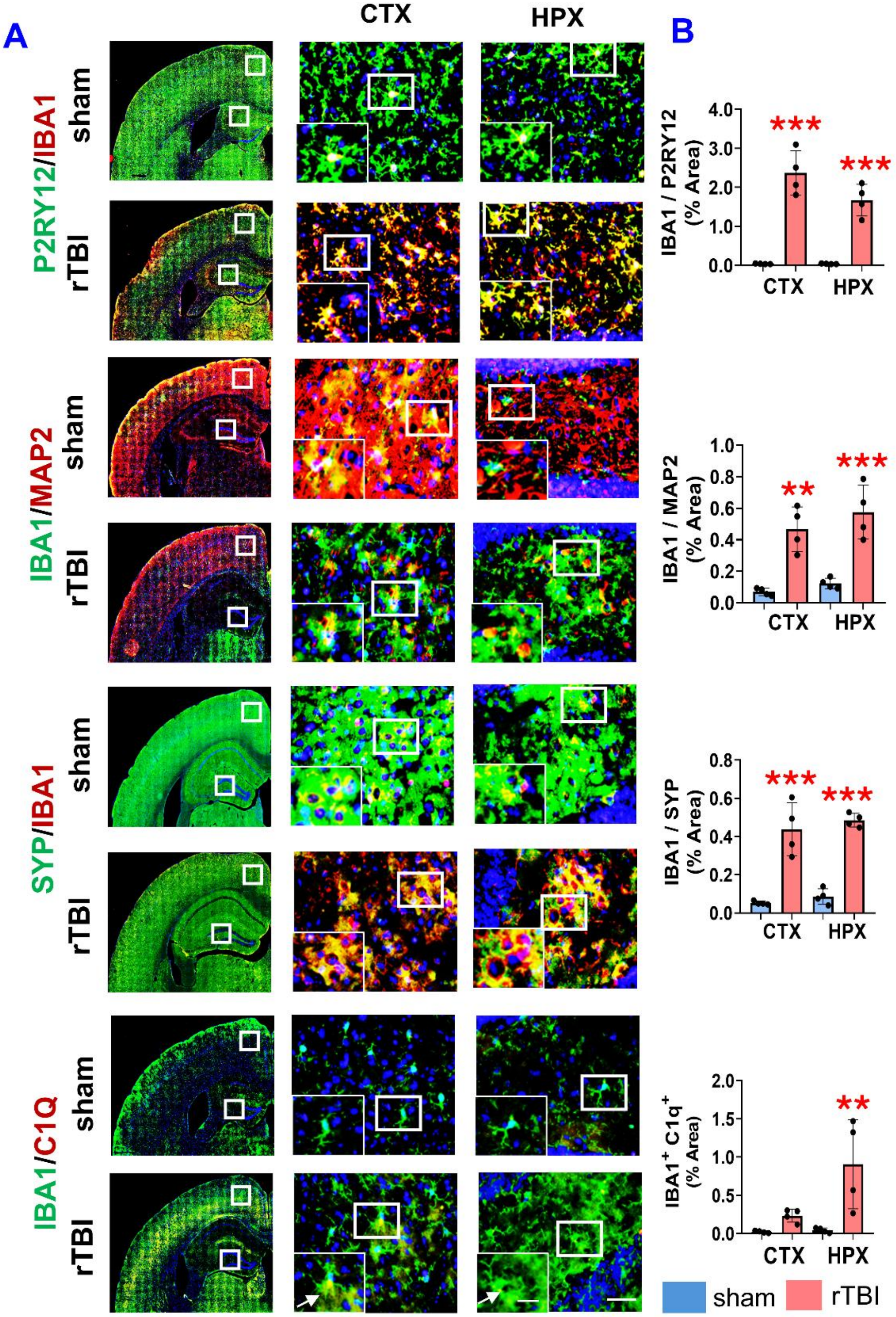

**Figure S6. rTBI induces chronic microglial activation and associated synaptic and neuronal damage at 120 dpi in rTBI mice. (A)** Representative immunofluorescence images showing double immunostaining of P2RY12 (Green, first panel), MAP2 (Red, second panel), SYP (Green, third panel), and C1q (Red, last panel) with IBA1 in the cortex (CTX) and hippocampus (HPX) at 120 dpi, illustrating microglial activation, complement engagement, and associated synaptic and neuronal alterations. **(B)** Histogram shows the quantification of the ratio of mean fluorescence intensity of IBA1/ P2RY12, IBA1/ MAP2, IBA1/ SYP, and IBA1+ C1q+ cells (%Area). Data are presented as mean ± SD. Scale bar: 50 µm (inset: 10 µm). *$p < 0.05$, **$p<0.01$, ***$p<0.001$ vs. sham; one-way ANOVA with Holm–Sidak test, n = 4/group.

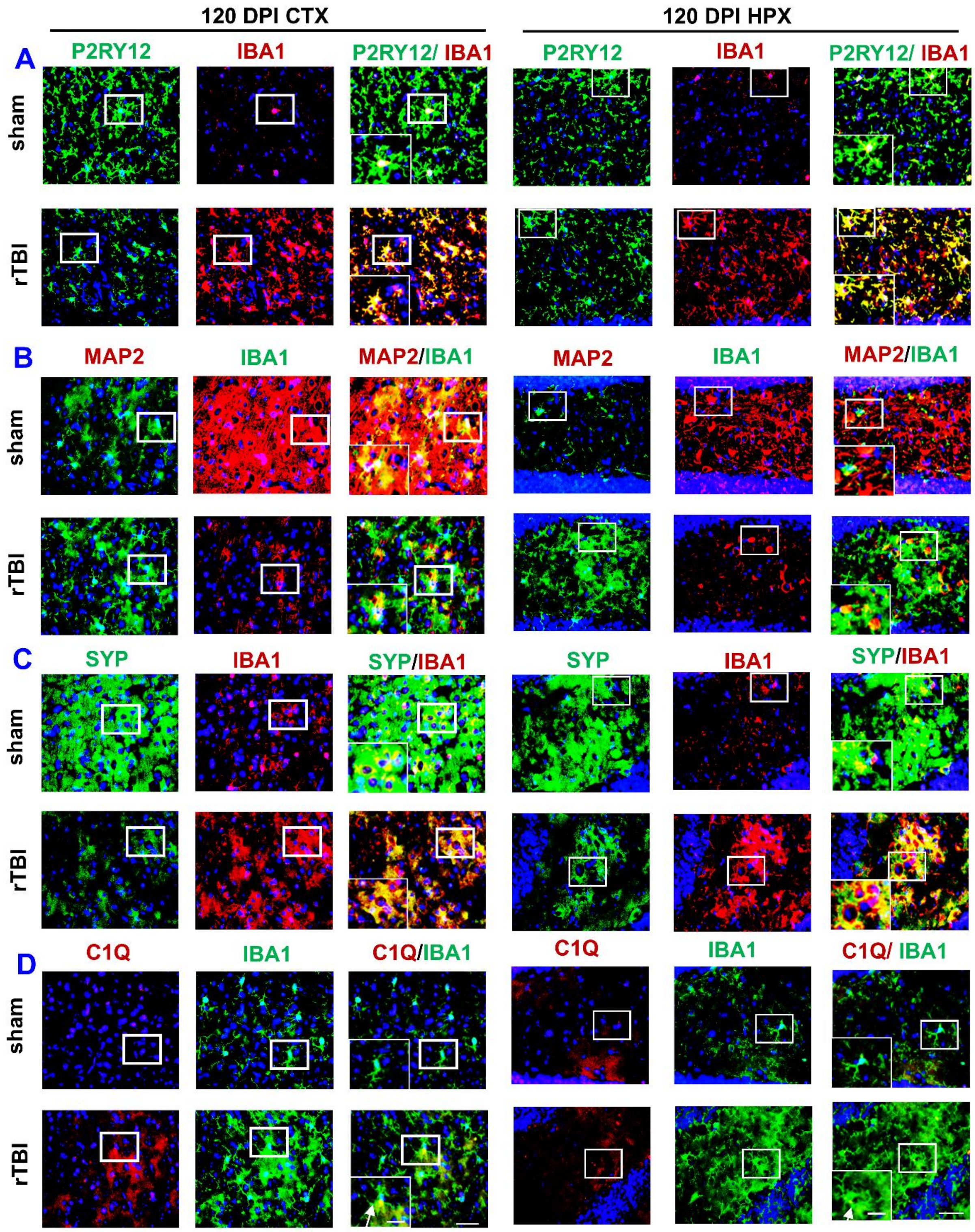

**Figure S7. rTBI induces chronic microglial activation and associated synaptic and neuronal damage at 120 dpi in rTBI mice.** Representative immunofluorescence images with individual channels showing double immunostaining of **(A)** P2RY12 (Green) / IBA1 (Red); **(B)** IBA1 (Green) / MAP2 (Red); **(C)** SYP (Green) / IBA1 (Red); **(D)** IBA1 (Green) and C1Q (Red), counterstained with DAPI in the cortex (CTX) and hippocampus (HPX) at 30 DPI

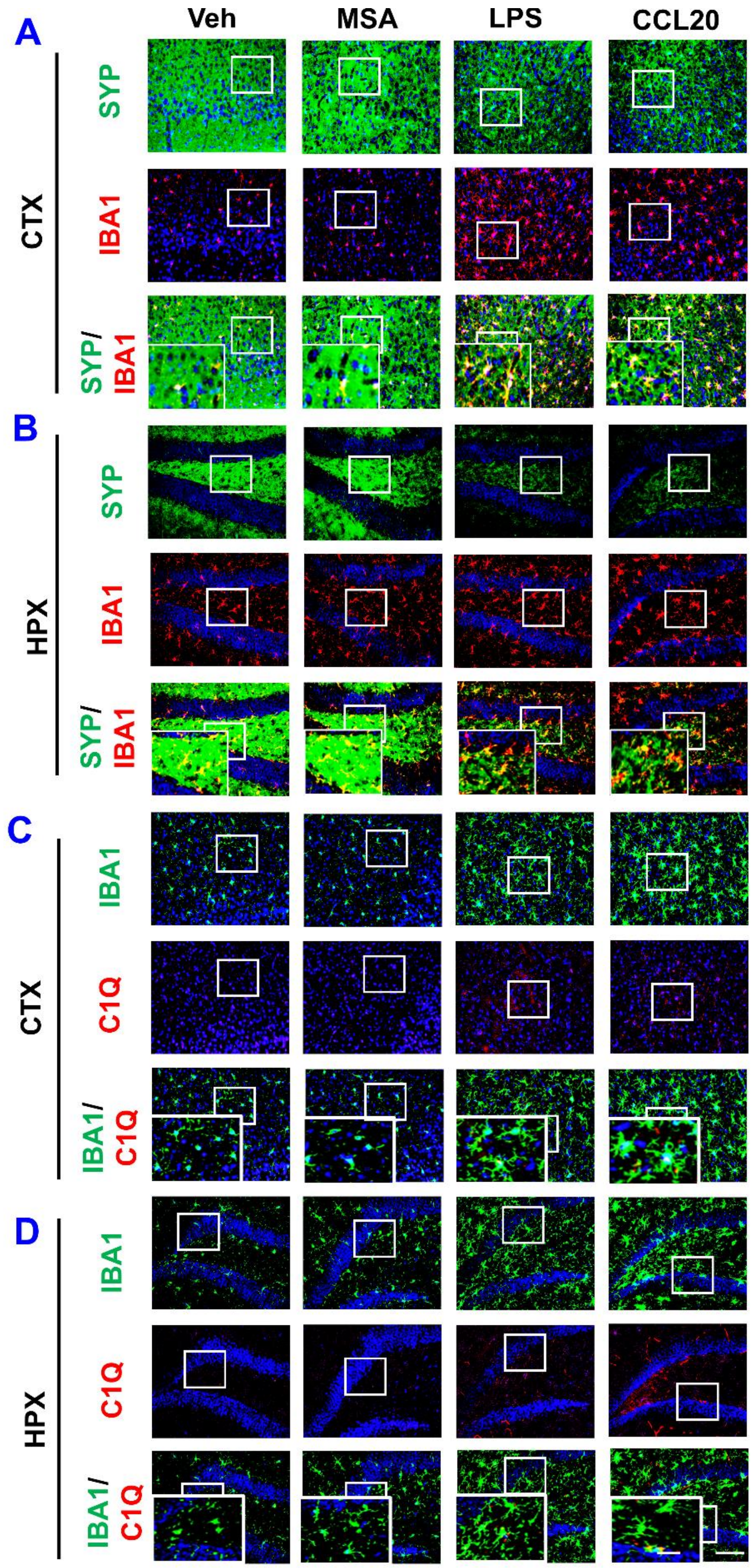

**Figure S8. CCL20 protein induces microglial activation and synaptic loss in C57BL/6 mice.** Representative immunofluorescence images with individual channels showing double labeling of **(A-B )** SYP (green) with IBA1 (red) and **(C-D)** IBA1 (green) with C1Q(red), counterstained with DAPI in the cortex (CTX) and hippocampus (HPX) at 14 days post-injection with Veh or Mouse Serum Albumin (MSA) or Lipopolysaccharide (LPS) or CCL20 recombinant protein

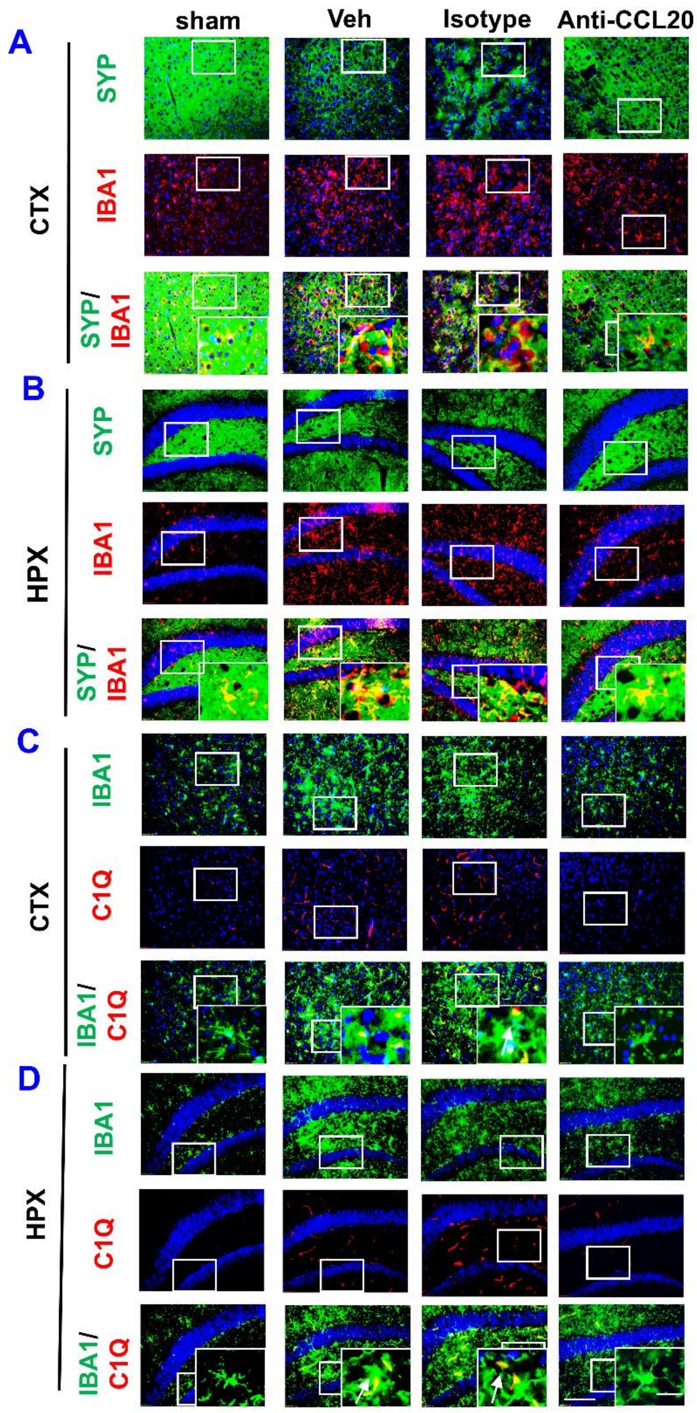

**Figure S9 Treatment with CCL20- neutralization antibody suppresses complement activation and synaptic pruning in rTBI mice.** Representative immunofluorescence images with individual channels showing double immunostaining of **(A-B)** SYP (green) with IBA1 (red) and **(C-D)** IBA1 (green) with C1Q(red), counterstained with DAPI in the cortex (CTX) and hippocampus (HPX) at 30 DPI.

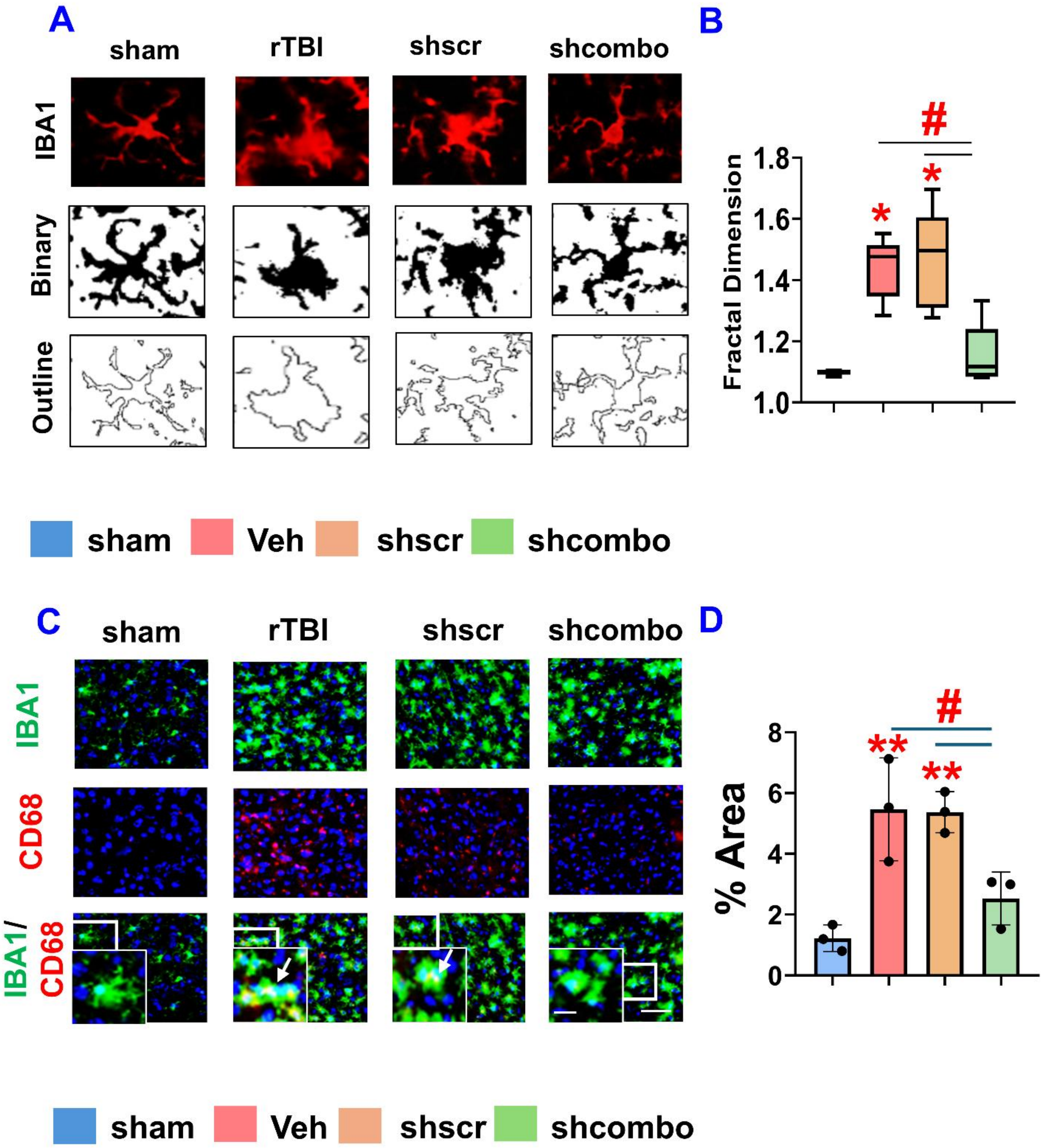

**Figure S10. Treatment with shCombo-DPX attenuates activated microglial phenotype in rTBI mice. (A)** Original and outlined microglia subjected to fractal analysis at 30 dpi. **(B)** Box plot showing fractal dimension of microglial phenotype in cortical regions of sham versus rTBI mice, n = 5/group. **(C)** Representative immunofluorescence images showing double labeling of CD68 (red) with IBA1 (green) in the cortex at 30 dpi, counterstained with DAPI. **(D)** Quantification of CD68 puncta localized to IBA1$^+$ microglia, expressed as % area. Data are presented as mean ± SD. Statistical analysis: one-way ANOVA with Holm–Sidak test; *$p < 0.05$, **$p < 0.01$, ***$p < 0.001$ vs. sham; #$p < 0.05$, ##$p < 0.01$ vs. treated groups.

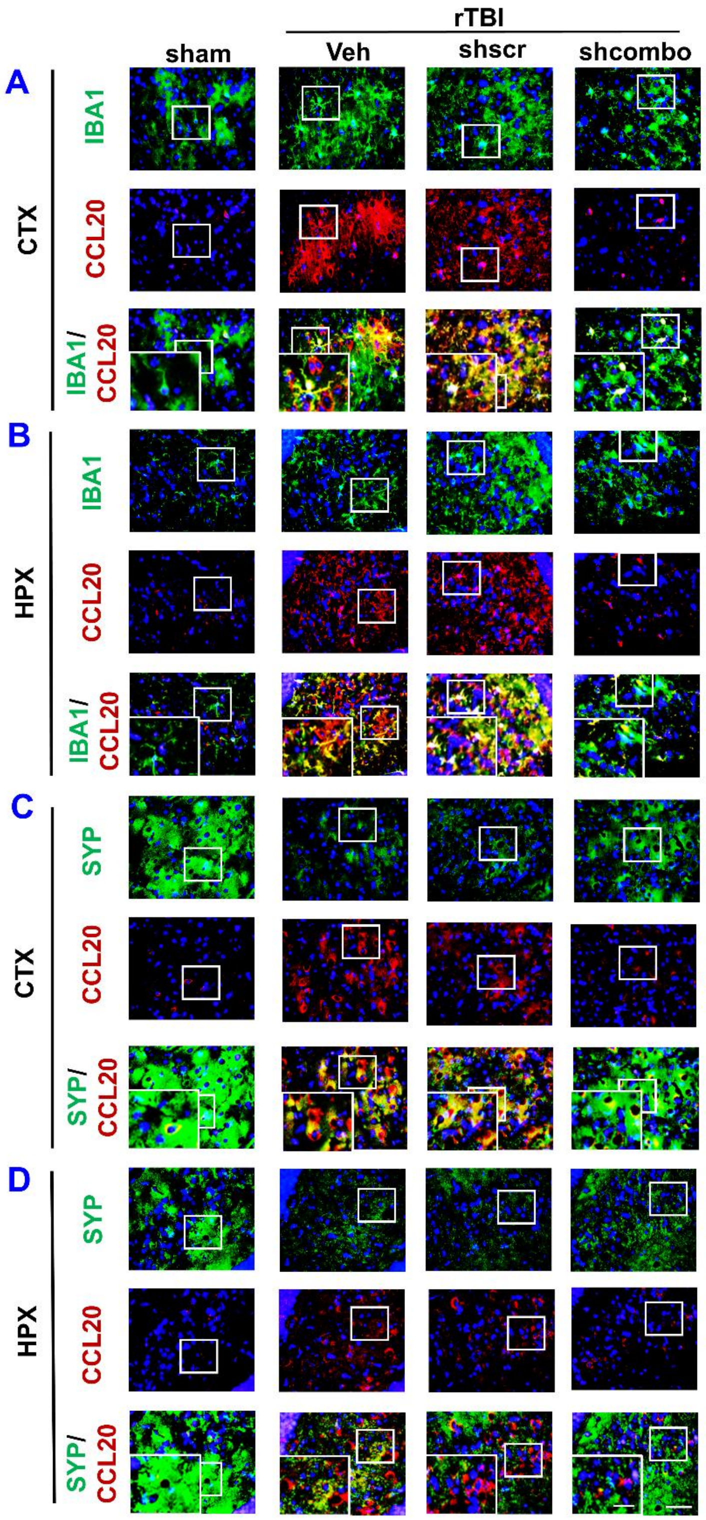

**Figure S11 Downregulation of the CCL20–CCR6 axis suppresses complement activation and synaptic pruning in rTBI mice .** Representative immunofluorescence images, individual channels showing double immunostaining of **(A-B)** IBA1 (Green) / CCL20 (Red) and **(C-D)** SYP (Green) / CCL20 (Red), counterstained with DAPI in the cortex (CTX) and hippocampus (HPX) at 30 DPI.

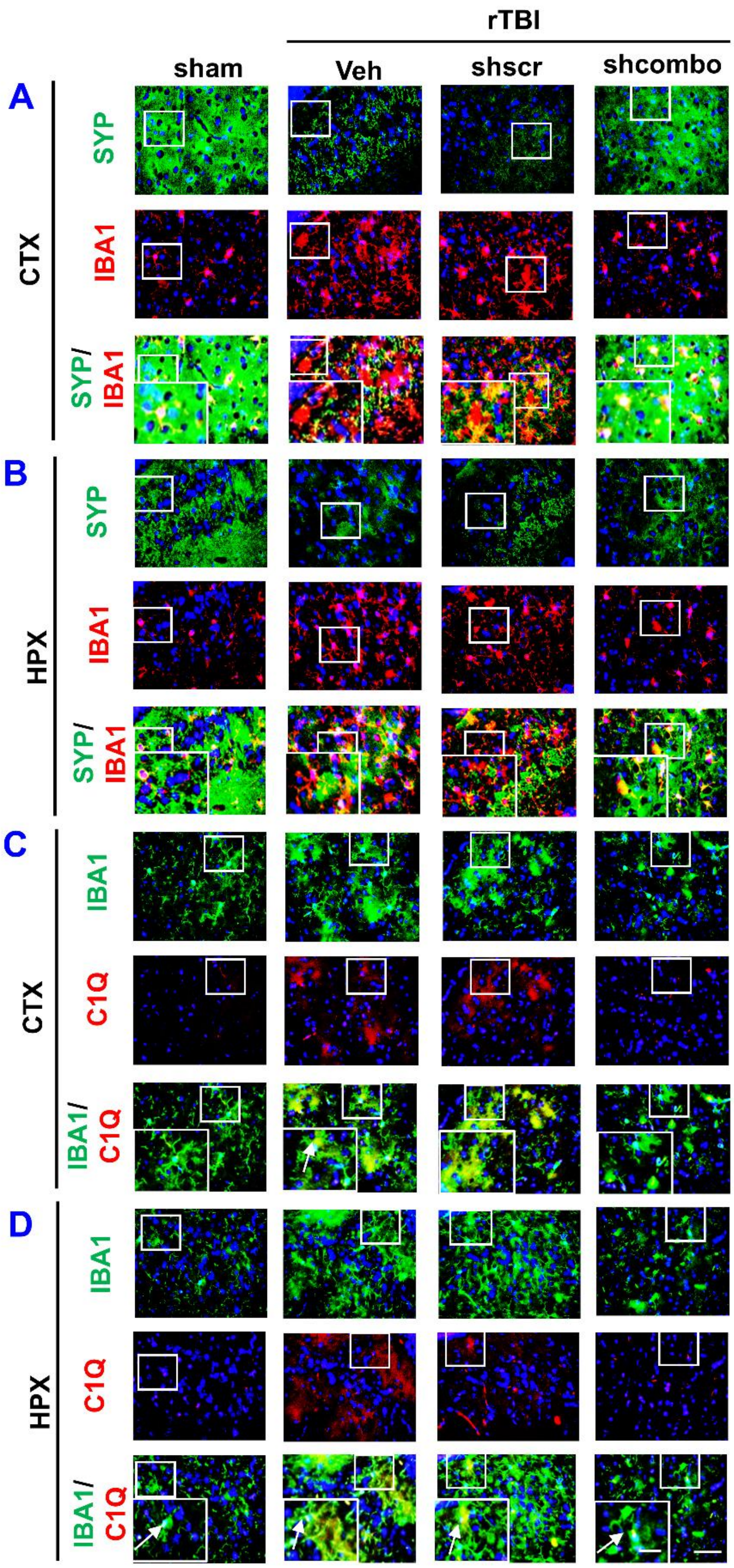

**Figure S12**. **Downregulation of the CCL20–CCR6 axis suppresses complement activation and synaptic pruning in rTBI mice .** Representative immunofluorescence images in individual channels showing double immunostaining of **(A-B)** SYP (Green) / IBA1 (Red); **(C-D)** IBA1 (Green) and C1Q (Red), counterstained with DAPI in the cortex (CTX) and hippocampus (HPX) at 30 DPI.

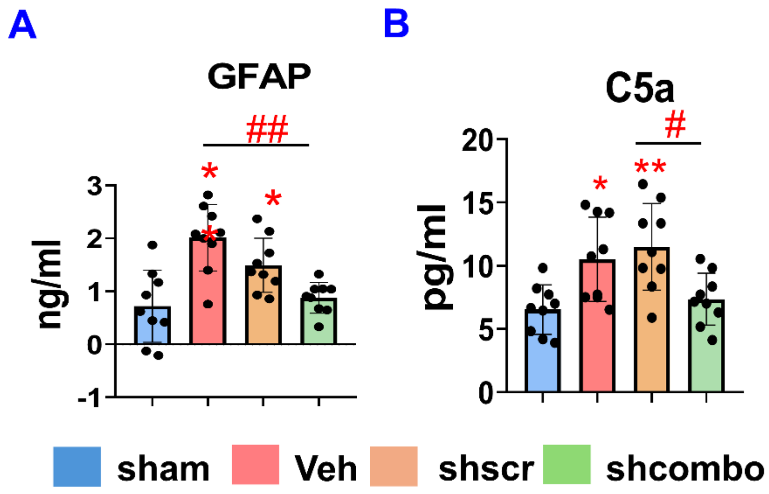

**Figure S.13 Treatment with shCombo-DPX attenuates peripheral inflammatory markers.** Circulating levels of GFAP **(A)** and C5a **(B)** measured in serum by ELISA. Data are presented as mean ± SD. Statistical analysis: one-way ANOVA with Holm–Sidak test; $*p < 0.05$, $**p < 0.01$, $***p < 0.001$ vs. sham; $\#p < 0.05$, vs treated groups.

### Open Field Test

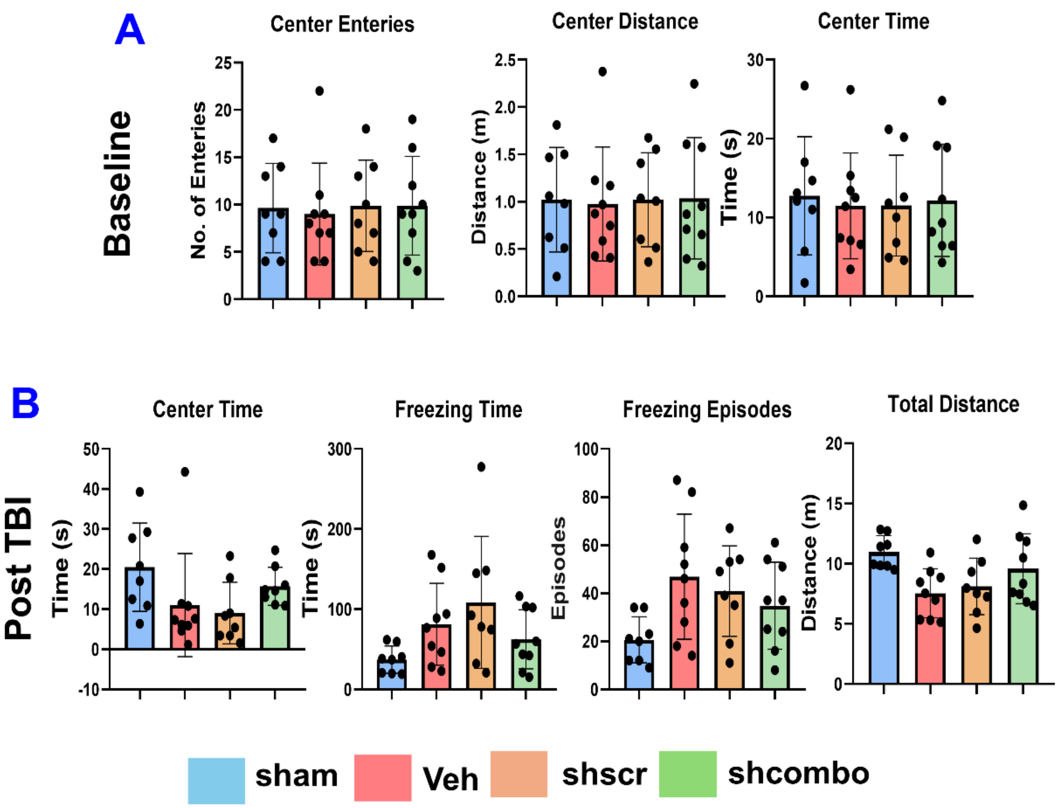

**Figure S14. (A-B) Behavioral outcomes in the open-field test 30 days after rTBI. (A)** Baseline changes prior to rTBI induction. **(B)** Center zone, total freezing time, freezing episodes, and total distance traveled in the open field

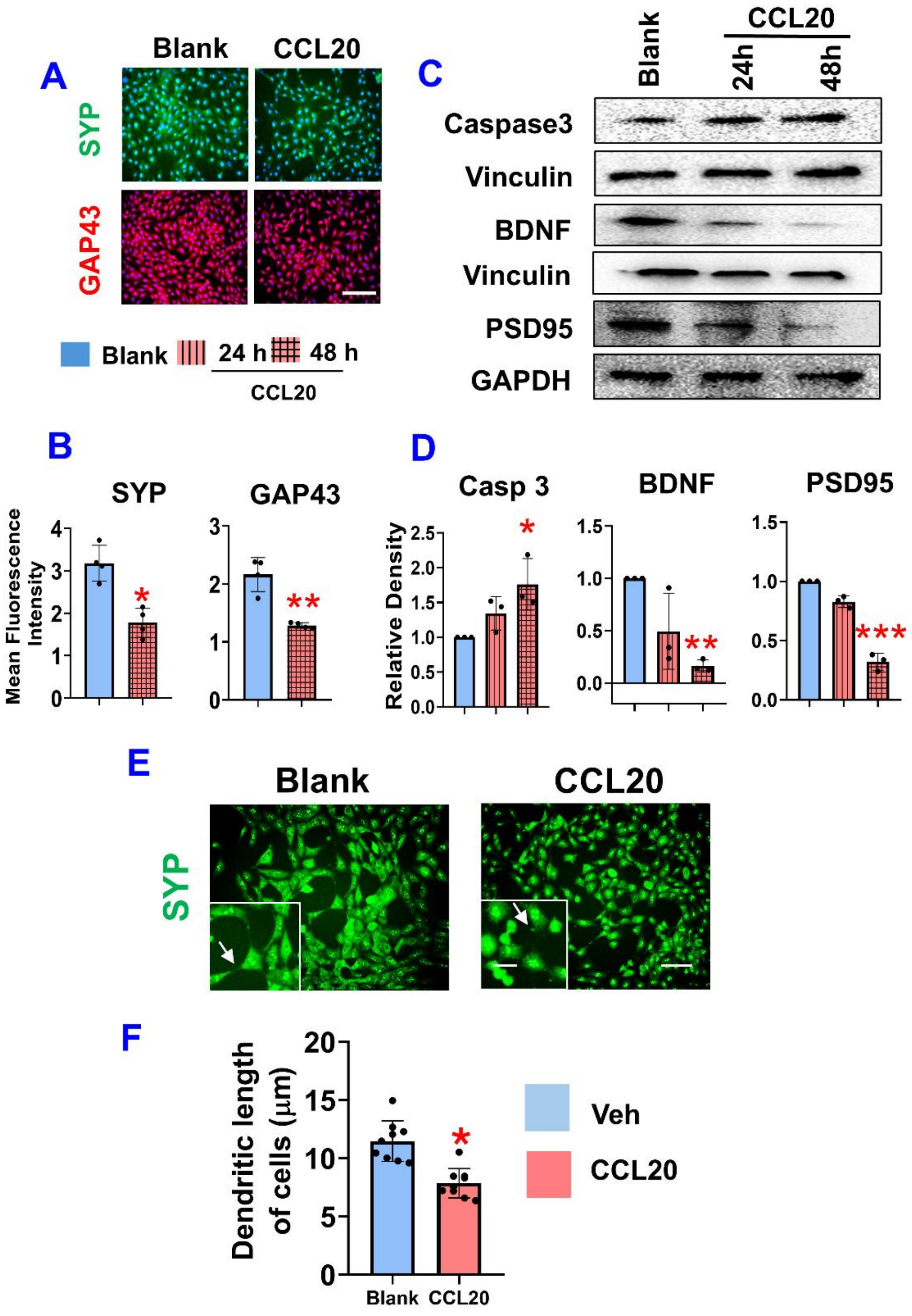

**Figure S15. CCL20 protein induces apoptosis and synaptic damage in HT22 neuronal (A)** Representative immunofluorescence images showing SYP and GAP43 staining in HT22 cells stimulated with CCL20 protein (500 ng/mL) for 48 h. **(B)** Quantification of SYP and GAP43 mean fluorescence intensity using ImageJ. **(C)** Representative immunoblots of apoptotic protein (Caspase-3) and synaptic plasticity markers (SYP, BDNF) in HT22 cells treated with CCL20 for 24 and 48 h. **(D)** Densitometric analysis of Caspase-3 and synaptic plasticity markers SYP and BDNF, normalized to vinculin or GAPDH. n=3/ group. Scale bar: 50 μm (inset: 10 μm). Data are presented as mean ± SD. *$p < 0.05$, **$p<0.01$, ***$p<0.001$ vs blank; $^{\#}$ $p < 0.05$, $^{\#\#}p < 0.01$, compared to treated groups; one-way ANOVA with Holm–Sidak test. **(E)** Representative immunofluorescence images showing synaptophysin staining. **(F)** Quantification of dendritic length (μm).

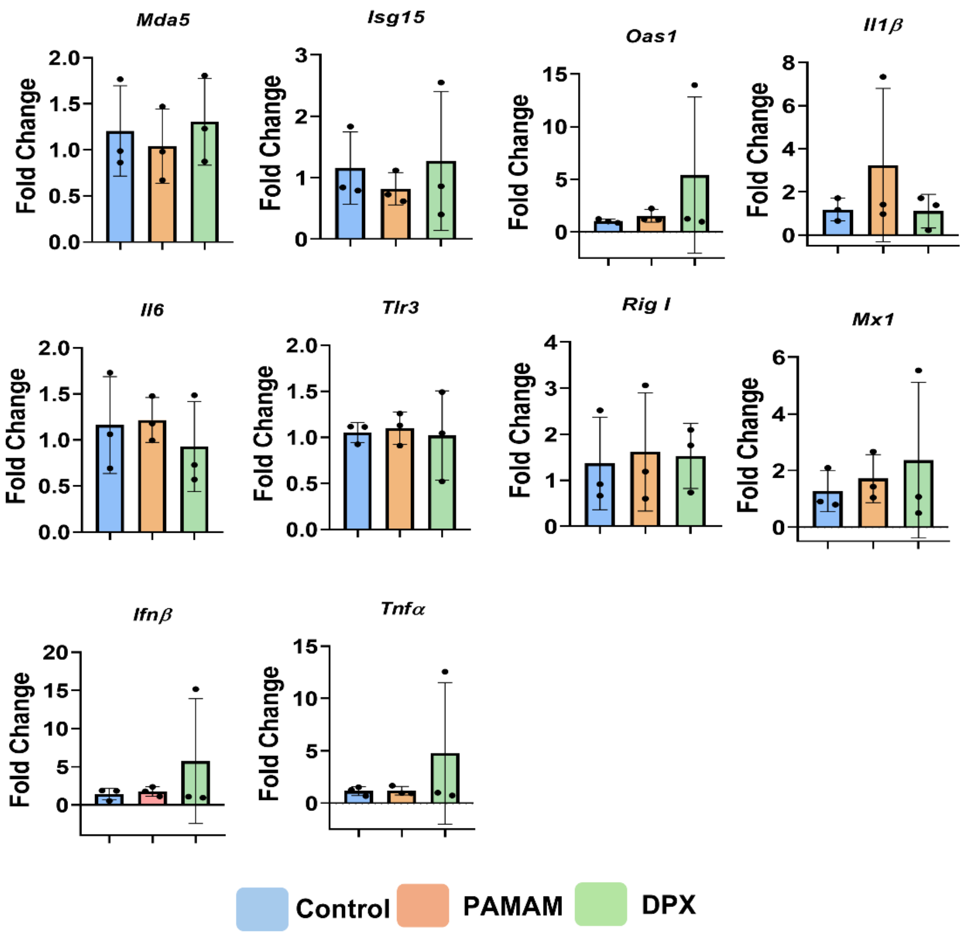

**Figure S16. PAMAM or shcombo DPX administration does not trigger immune responses in mice.** Histograms showing gene expression of *Mda5, Tlr3, Isg15, Rig-I, Oas1, Mx1, Ifnβ, Il6, Il1β, and Tnfα* assessed by qPCR from RNA isolated from cortical tissue (CTX) at 30 dpi.

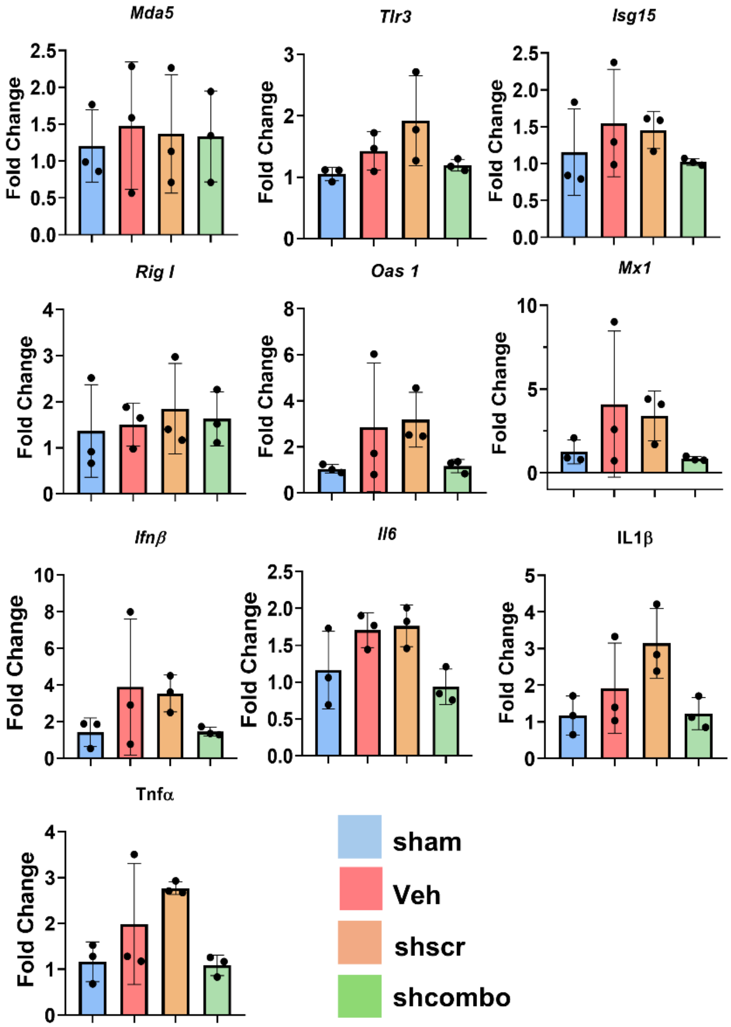

**Figure S17. Shcombo DPX administration does not trigger immune responses in rTBI mice.** Histograms showing gene expression of *Mda5, Tlr3, Isg15, Rig-I, Oas1, Mx1, Ifnβ, Il6, Il1β, and Tnfα* assessed by qPCR from RNA isolated from cortical tissue (CTX) at 30 dpi.

Supplemental Table

**Table S.1. Ingenuity Pathway Analysis (IPA) illustrating observed and predicted CCL20-centered interaction networks in the cortex.**

| | | | | | | |
|---|---|---|---|---|---|---|
| **DEP Connected to CCL20** | | | | | | |
| ACVR1 | ACVR2A | BAG3 | BAG6 | DBN1 | CANX | ADCK2 |
| ACIN1 | DBNL | APLP2 | CFDP1 | ACLY | CLU | DKC1 |
| HSPA8 | BCAR1 | AHSG | CREBBP | EEF1G | DAB2 | EPB41L1 |
| GLS | G3BP2 | DDX21 | LRRC59 | DEGS1 | DEPTOR | HDAC4 |
| KRAS | COL4A2 | F3 | ASL | EN01 | DZIP3 | MYBBP1A |
| KRT8 | FGA | FLNC | LMNA | LAMP1 | MYL12B | LYAR |
| PANPC1 | NCAM1 | LPIN1 | PSPC1 | MAPK8 | METTL3 | MED6 |
| ITGB8 | KMT2A | PLS3 | MYH9 | PPM1A | NLGN3 | PURB |
| NSRP1 | ICAM1 | MAGI1 | PLD2 | PEBP1 | PCOLCE | PKM |
| RAB3GAP1 | PAR | RMC1 | PRKCQ | PPP1R10 | RPF2 | MYO5A |
| PURA | APOB | PRKAB1 | S100A6 | STK25 | PXMP2 | RAB227A |
| SARS1 | SF3B2 | TBL1XR1 | TPM1 | SRP19 | SPTBN1 | SLC5A5 |
| SQOR | S100A4 | RPL18A | APOA1 | SERBP1 | CTSD | SMAD3 |
| CD14 | STXBP1 | VCP | YWHAQ | MTDH | ZC3H4 | SMYD3 |
| TNC | USP7 | SELENBP1 | SELENBP2 | | | |
| **Proteins linking CCL20 to Complement** | | | | | | |
| STING1 | BMI1 | DOS | STAT6 | PPARD | OCLN | IRF4 |
| CCR6 | IRF1 | TP53 | SP1 | BCL3 | RXPA | NFKB2 |
| NFKB1 | STAT3 | GPCR | RXRB | NRF1 | RELA | JUN |
| NFKBIA | YBX1 | CEBPA | AHR | ESR2 | JUNB | ESR1 |
| LGALS3 | USO1 | CEBPB | SMARCC1 | FOXO1 | SMAD2 | MEOX2 |
| HMGB1 | NFE2L2 | APC | POU5F1 | RELS0X2 | WWOX | IFI16 |
| STAT1 | NR3C1 | E2F3 | NR2F2 | DDX5 | | |
| **Complement Proteins** | | | | | | |
| C1QBP | C1QB | C3 | C1OL1 | C5 | C1QC | |
| **DEP connected to Complement** | | | | | | |
| GFAP | S100A10 | DCTN1 | ABI3 | DDX19B | DYNC1H1 | DYNC1LI2 |
| DYNLL2 | CTSS | HMGCLL1 | GBP2 | NRXN3 | AP3D1 | VAV1 |
| KLHL7 | POLR2M | GSK3A | NLGN2 | SLC4A1 | PSMC1 | YIF1B |
| YIPF3 | MAP3K4 | UBR4 | | | | |

**Table S.2. Ingenuity Pathway Analysis (IPA) illustrating observed and predicted CCL20-centered interaction networks in the Hippocampus.**

| DEP Connected to CCL20 | | | | | | |
|---|---|---|---|---|---|---|
| GOLM1 | ESRRA | S100A5 | MAP4K1 | | | |
| **Proteins linking CCL20 to Complement** | | | | | | |
| IGFBP5 | APOE | STAT3 | CTSD | PLAT | CDK4 | PTPRC |
| CD68 | ACTB | PEBP1 | ITGB2 | LY86 | B2M | STING1 |
| **Complement Proteins** | | | | | | |
| C1QC | C1QA | C3 | C4A/B | C5 | C2 | C6 |
| **DEP connected to Complement** | | | | | | |
| CCNE1 | SFXN4 | ZNF106 | RPS21 | RPS29 | RHOB | MANBA |
| UCHL1 | CD55 | SERPINA 3N | CD63 | | | |

**Table S.3. Serum liver enzyme levels (ALT, AST) in Mouse**

| | ALT(U/L) | AST(U/L) |
|---|---|---|
| | Normal Range (28-132) | Normal Range (59-247) |
| Sham1 | 49 | 49 |
| Sham2 | 70 | 97 |
| shScr 1 | 44 | 52 |
| shScr 2 | 196 | 301 |
| shScr 3 | 32 | 46 |
| shCombo 1 | 32 | 43 |
| shCombo 2 | 33 | 46 |
| shCombo 3 | 27 | 65 |

**Table S 4 showing complete blood count (CBC) panel, presented alongside established normal ranges**

| | WBC (k/ul)) | LYM (k/ul) | NEUT(k/ul) | MONO (k/ul) | EOS/GRAN(k/ul) |
|---|---|---|---|---|---|
| | 5-10 | 4.5-8.9 | 0.50-0.70 | 0.3-0.7 | 0.0-0.34 |
| Sham1 | 8.88 | 7.95 | 0.6 | 0.17 | 0.15 |
| Sham2 | | | | | |
| shScr 1 | 13.51 | 12.22 | 0.54 | 0.61 | 0.14 |
| shScr 2 | 6.36 | 5.48 | 0.45 | 0.34 | 0.08 |
| shScr 3 | 9.24 | 7.49 | 1.18 | 0.44 | 0.12 |
| shCombo 1 | 8 | 7.25 | 0.37 | 0.29 | 0.08 |
| shCombo 2 | 12.29 | 11.36 | 0.54 | 0.23 | 0.14 |
| shCombo 3 | 2.98 | 2.64 | 0.2 | 0.1 | 0.04 |

| | BASO/GRAN(k/ul) | RBC(M/ul) | MCV (fL) | HGB (g/dL) | PLT(M/ul) | MPV(fL) |
|---|---|---|---|---|---|---|
| | 0.00-0.08 | 8.7-10.1 | 53.6-57.2 | 14.4-16.8 | 855-1911 | 4.6-5.5 |
| Sham1 | 0.01 | 9.6 | 53.6 | 14.8 | 734 | 7 |
| Sham2 | | 9.89 | 53.6 | 15 | 458 | 7.1 |
| shScr 1 | 0 | 10.48 | 51.3 | 15.5 | 1182 | 7 |
| shScr 2 | 0.01 | 10.34 | 53.51 | 15.2 | 585 | 7.4 |
| shScr 3 | 0.01 | 10.29 | 53.6 | 15.5 | 961 | 6.6 |
| shCombo 1 | 0.01 | 8.82 | 52.3 | 13.2 | 665 | 6.9 |
| shCombo 2 | 0.02 | 10.18 | 52.8 | 15.3 | 1069 | 6.4 |
| shCombo 3 | 0 | 6.9 | 50.6 | 10.6 | 284 | 6.9 |

**Table S5 shCCL20 and shCCR6 target sequence:**

| shCCL20 target sequence | shCCR6 target sequence | shScramble sequence |
|---|---|---|
| GGCCGATGAAGCTTGTGACATT | GAACTGCCCACTTCCCTTTCTA | ACCTAAGGTTAAGTCGCCCTCG, |
| GGCGCTTAGTGGAAGGATTAAT | GATCCATGACTGACGTCTACCT | ACCTAAGGTTAAGTCGCCCTCG |
| AGAGCTATTGTGGGTTTCA | GCTGCAGGATCGTGATGTC | ACCTAAGGTTAAGTCGCCC, |
| CGTACATACAGACGCCTCT | GTATCAGCATGGACCGGTA | ACCTAAGGTTAAGTCGCCC |

**Table S6. Forward and reverse primer sequences used for qPCR.**

| Gene | Forward Primer | Reverse Primer |
|---|---|---|
| CCL20 | 5`-ATGGCCTGCGGTGGCAAGCGTCTG-3 | 5`-TAGGCTGAGGAGGTTCACAGCCCT-3` |
| CCR6 | 5`-CCTCACATTCTTAGGACTGGAGC-3` | 5`-GGCAATCAGAGC TCTCGGA-3` |
| IL6 | 5`-TACCACTTCACAAGTCGGAGGC-3` | 5`-CTGCAAGTGCATCATCGTTGTTC-3` |
| IFNB1 | 5`-GCCTTTGCCATCCAAGAGATGC-3` | 5`-ACACTGTCTGCTGGTGGAGTTC-3` |
| ISG15 | 5`-CATCCTGGTGAGGAACGAAAGG-3` | 5`-CTCAGCCAGAACTGGTCTTCGT-3` |
| OAS1 | 5`-GAGGTGGAGTTTGATGTGCTGC-3` | 5`-GTGAAGCAGGTAGAGAACTCGC-3` |
| MX1 | 5`-TGGACATTGCTACCACAGAGGC-3` | 5`-TTGCCTTCAGCACCTCTGTCCA-3` |
| TLR 3 | 5`-GTCTTCTGCACGAACCTGACAG-3` | 5`-TGGAGGTTCTCCAGTTGGACCC-3` |
| IL1β | 5`- TGGACCTTCCAGGATGAGGACA-3` | 5`-GTTCATCTCGGAGCCTGTAGTG-3` |
| RIGI | 5`-AGCCAAGGATGTCTCCGAGGAA-3` | 5`-ACACTGAGCACGCTTTGTGGAC-3` |
| MDA5 | 5`-TGCGGAAGTTGGAGTCAAAGCG-3` | 5`-CACCGTCGTAGCGATAAGCAGA-3` |
| BETA ACTIN | 5` -GTATGCCTCGGTCGTACCA-3` | 5` -CTTCTGCATCCTGTCAGCAA-3` |

### Table S7 Primary and secondary antibodies used for immunostaining.

| Primary antibody | Source | Catalog No. | Dilution | Secondary antibody | Source | Catalog No. | Dilution | Development |
|---|---|---|---|---|---|---|---|---|
| Rabbit anti-IBA1 | Wako | 019-19741 | 1:500 | Alexafluor 594- anti rabbit/ Alexafluor 488- anti rabbit | Invitrogen/ Invitrogen | A11012/ A21206 | 1:1000 | Fluorescence/ Fluorescence |
| Chicken anti-GFAP | Millipore Sigma | AB5541 | 1:1000 | Alexafluor 594- anti chicken/ Alexafluor 488- anti chicken | Invitrogen/ Invitrogen | A11042/ A11039 | 1:1000 | Fluorescence/ Fluorescence |
| Rabbit anti-CCR6 | Abcam | Ab78429 | 1:500 | Alexafluor 594- anti rabbit | Invitrogen | A11012 | 1:1000 | Fluorescence |
| Rabbit anti-CCL20 | Abcam | Ab9829 | 1:500 | Alexafluor 594- anti rabbit/ Alexafluor 488- anti rabbit/ Biotinylated goat anti-rabbit | Invitrogen/ Invitrogen/ Vector Laboratories Inc., Burlingame, Ca. | A11012/ A11008/ BA-1000 | 1:1000/ 1:1000/ 1:400 | Fluorescence/ Fluorescence/ DAB |
| Mouse anti-NeuN | Abcam | ab104224 | 1:1000 | Alexafluor 594- anti mouse | Invitrogen | A21203 | 1:1000 | Fluorescence |
| Rabbit anti-P2Y12 | Abcam | Ab300140 | 1:1000 | Alexafluor 488- anti rabbit | Invitrogen | A21206 | 1:1000 | Fluorescence |
| Rabbit anti-MAP2 | Abcam | Ab183830 | 1:1000 | Alexafluor 594- anti rabbit | Invitrogen | A11012 | 1:1000 | Fluorescence |
| Rabbit anti-SYP | Proteintech | 17785-1-AP | 1:1000 | Alexafluor 594- anti rabbit/ Alexafluor | Invitrogen/ Invitrogen | A11012/ A21206 | 1:1000 | Fluorescence |

**Table S.8. Primary and secondary antibodies used for Western blotting.**

| Primary antibody | Source | Catalog No. | Dilution | Secondary antibody | Source | Catalog No. | Dilution | Development |
|---|---|---|---|---|---|---|---|---|
| Caspase-3 | Cell signaling | 14220T | 1:1000 | Anti- rabbit IgG HRP | Invitrogen | 31460 | 1:1000 | HRP |
| BDNF | Abcam | Ab108319 | 1:1000 | Anti- rabbit IgG HRP | Invitrogen | 31460 | 1:1000 | HRP |
| PSD95 | Cell signaling | 2507 | 1:1000 | Anti- rabbit IgG HRP | Invitrogen | 31460 | 1:1000 | HRP |
| Vinculin | Cell signaling | 13901 | 1:1000 | Anti- mouse IgG/IgM HRP | EMD Millipore Corp., USA | AP130P | 1:1000 | HRP |
| GAPDH | Cell signaling | 2118 | 1:1000 | Anti- rabbit IgG HRP | Invitrogen | 31460 | 1:1000 | HRP |
